# The *ift140*-Deficient Zebrafish: A Model for Renal Cystogenesis and an F0-Based Screen to Identify Genetic Modifiers of Kidney Cysts

**DOI:** 10.1101/2025.01.02.631132

**Authors:** Ping Zhu, Andrew Lavin, Yonghe Ding, Xiaolei Xu, Xueying Lin

## Abstract

Genetic modifiers are believed to play an important role in the onset and severity of polycystic kidney disease (PKD), but identifying these modifiers has been challenging due to the lack of effective methodologies. In this study, we investigated zebrafish mutants of *IFT140*, a newly identified ADPKD gene, and observed phenotypes similar to those seen in mammalian models, including kidney cysts and bone defects. Using efficient microhomology-mediated end joining (MMEJ)-based genome editing technology, we demonstrated that *ift140* CRISPRants recapitulate mutant phenotypes while bypassing the early lethality of the mutants to allow for renal cyst analysis in adult fish. In addition to cilia defects, non-cilia phenotypes, such as disrupted cell polarity and aberrant microtubule stabilization in kidney epithelial cells, may also contribute to *ift140*-associated cystogenesis. Importantly, the ability to detect *ift140*-associated renal cysts with ease allowed us to develop an F0-based genetic screen to identify potential protective modifiers. A pilot screen of sixteen genes previously linked to dysregulated signaling pathways in ADPKD revealed both known and novel modifiers, including *mtor* and *ulk1a*. Inhibition of *mtor* and *ulk1a* reversed both cilia-related and non-cilia-related abnormalities. In summary, our study underscores the potential of using zebrafish as a model for the efficient discovery of genetic modifiers of kidney cysts.

## Introduction

Autosomal dominant polycystic kidney disease (ADPKD) is one of the most prevalent and potentially lethal genetic diseases. Many patients eventually develop renal failure and require dialysis or a kidney transplant^1–4^. High phenotypic variations have been noted in ADPKD patients. While many factors contribute to the variations, the intrafamilial phenotype differences strongly suggest the involvement of modifier genes^5–8^. Genome-wide association studies (GWAS) in human populations and quantitative trait locus (QTL) analysis in rodents are ongoing methods for the identification of genetic modifiers. Based on co-inheritance of mutations in certain genes together with *PKD1* or *PKD2*, a limited number of modifier genes such as *HNF1β*, *TSC2*, and *DKK3* have been reported^9–11^. However, these statistics-based methods have significant shortcomings: 1) a precise genotype-phenotype relationship is difficult to establish and 2) experimental validation of a candidate modifier in rodent models has very low-throughput, requiring mutants and multi-generation crosses. Consequently, the identity of ADPKD genetic modifiers remains largely unknown, which severely hinders the development of therapeutic intervention. Thus, an alternative animal model with more efficient genetic tools and a novel method enabling rapid discovery of genetic modifiers of ADPKD are highly desirable.

Zebrafish have been used to study cystogenesis. The zebrafish pronephric kidney is simply composed of a pair of tubules that are fused at a glomerulus; mutations in genes associated with human cystic diseases, such as *pkd1* and *hnf1b*, have been found to cause kidney cysts in zebrafish embryos^12–14^. Large scale compound screens have been conducted in zebrafish embryos to search for therapeutic drugs for PKD, and some candidates showed promising effects in mammalian models^15,16^. These screens use characteristic body curvature of *pkd2* mutants as a surrogate because a kidney cyst-based assay cannot be conducted using currently available mutants of ADPKD genes. *pkd1* mutants do form pronephric cysts, featuring low penetrance at 2 days post fertilization (dpf), full penetrance at 3 dpf, and then complicated by the development of edema. Visual observation of cysts at 3 dpf is blocked from the dorsal view because the anterior pronephron is now located ventral of the head and is unreliable from the lateral view because the cyst is not big enough yet. Consequently, pronephric cysts can only be reliably detected at 3 dpf via section-HE staining^14^, which is not a suitable method of screening. On the other hand, *pkd2* mutants do not form pronephric cyst^13,17^. Nonetheless, large scale genetic screen, particularly genetic modifier screen has not been executed yet for any PKD models.

Recently, CRISPR-induced mosaic knockout has been used to identify genes implicated in morphogenesis, regeneration, and behavior phenotypes in the zebrafish embryos^18–21^. However, multi-locus targeting of a single gene is needed to achieve functional disruption due to the non-homologous recombination mechanism of CRISPR knockout^21^. On the other hand, microhomology-mediated end joining (MMEJ)-based CRISPR can achieve highly efficient knockout via single-locus targeting because frame- shifting genetic lesions can be predicted and pre-selected^22^. Using the MMEJ technology, genetic modifiers of cardiomyopathy have been successfully identified via F0-based screens both in zebrafish embryos and adults^23–25^.

*IFT140* encodes a core component of the intraflagellar transport (IFT) complex A. Biallelic mutation of *IFT140* is associated with skeletal ciliopathies, such as the short rib thoracic dysplasia (SRTD) and Mainzer-Saldino syndrome (MSS), with extraskeletal phenotypes including kidney cysts^26,27^. In experimental models, *Ift140* ablation causes skeletal ciliopathies, cystic kidneys, etc.^28,29^. Recently, *IFT140* was implicated as an important ADPKD gene, which for the first time directly links a ciliary structure gene to human cystic disease^30^. As a retrograde IFT, IFT140 coordinates the removal of cargos from cilia, it also regulates the entry of proteins into cilia^31–33^. However, little is known about the mechanisms underlying its functions in renal cyst development.

Here, we generated a zebrafish mutant of *IFT140*. Inactivation of *ift140* resulted in profound renal cysts and skeletal formation defects. In the kidney epithelial cells, single cilium was progressively shortened, multi-cilia bundles were disorganized, and cytoskeletal microtubular dynamics were altered. Furthermore, we showed that *ift140^MJ^*, a MMEJ-based CRISPRant, faithfully recapitulate pronephric cyst phenotype of the mutants, enabling a genetic screening strategy that directly based on the renal cyst phenotype. A pilot F0 genetic screen identified both known and novel genetic modifiers that protect against kidney cysts, which were later validated in stable mutants. Our study highlights the potential of using zebrafish *ift140* platform for large scale screen of genes that protect against kidney cysts.

## Results

### *ift140^-/-^* fish recapitulates renal and bone phenotypes in mammalian models

Considering that *IFT140* is the third most mutated gene in ADPKD patients^30^, we decided to investigate its function in zebrafish. Only one orthologue of mammalian *IFT140* has been identified in zebrafish, located on chromosome 24 (zfin.org). *In situ* hybridization revealed weak expression of the zebrafish *ift140* transcript in the kidney and other regions such as the notochord (Supplemental Figure 1). We generated a stable mutant of *ift140* (Figure 1A), which showed an absence of Ift140 protein expression (Figure 1B), suggesting a null mutant allele. All *ift140^e2/e2^* embryos developed distinctive pronephric cysts, detectable as early as 2 days post fertilization (dpf) (Figure 1C, 1D). Despite the progressively enlarged pronephric cysts, *ift140^e2/e2^* embryos exhibited generally normal gross morphology, without abnormal body curvature or signs of cardiac or whole-body edema that frequently found in other cilia mutants (Supplemental Figure 2A). However, at approximately 2 weeks of age, the mutant larvae were significantly smaller and began to die, with none surviving past 30 dpf (Supplemental Figure 2B, 2C).

**Figure 1.**
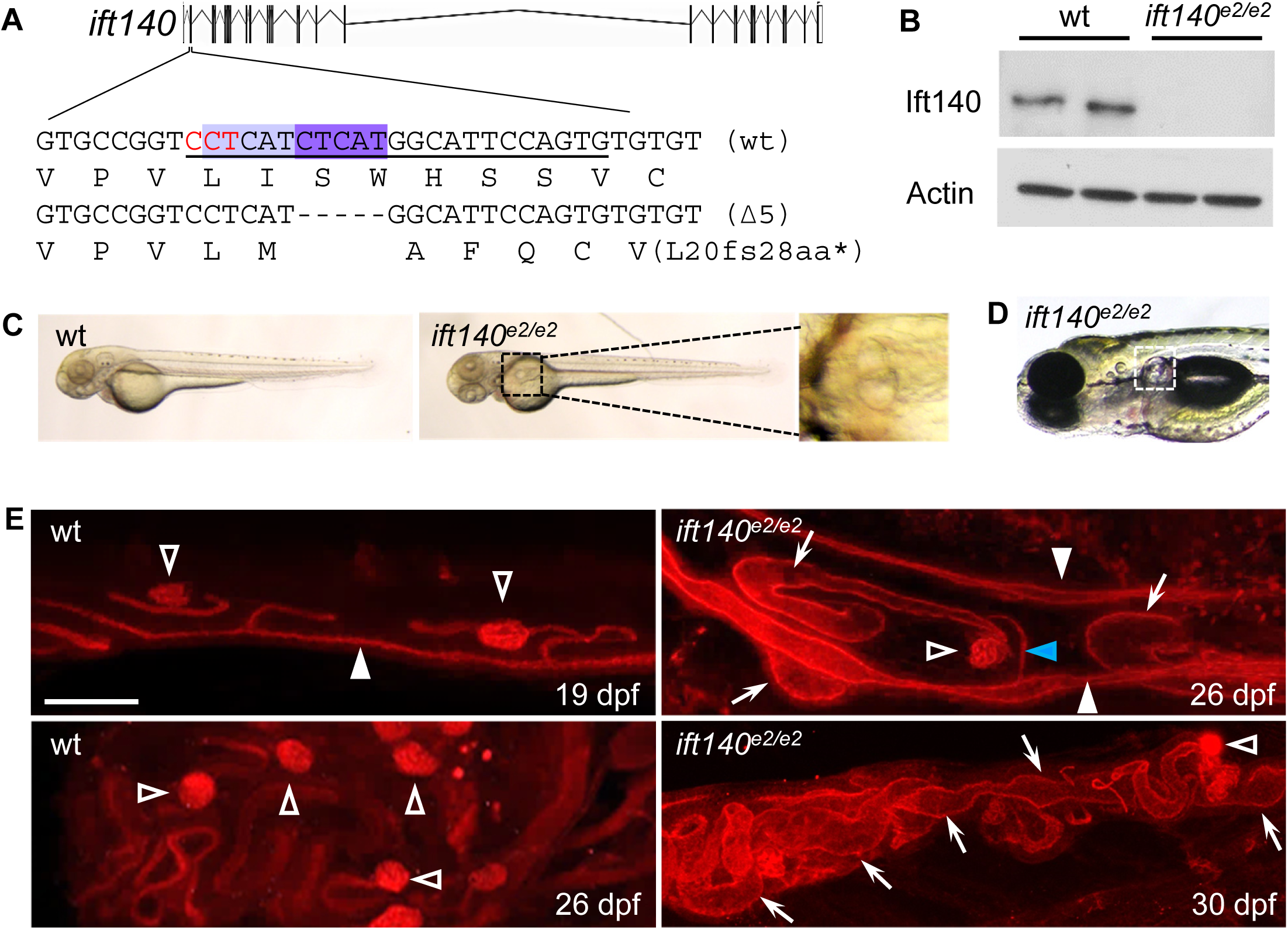
Inactivation of *ift140* results in the formation of kidney cysts in both the pronephros and the newly formed mesonephros. (**A**) Schematic representation of MMEJ-mediated genetic lesions in *ift140*. Underlined: *sgRNA* target sequence; red: Pam sequence; shaded: microhomology sequence. Nucleotide deletions (indicated by dashes) cause a coding frameshift that presumably results in a premature stop codon (*). The corresponding amino acid sequence is shown below the DNA sequence. (**B**) Ift140 protein expression is abolished in homozygous *ift140* mutants. Five wild-type and five mutant zebrafish were collected for western blot analysis at 14 dpf, and representative images are shown. (**C, D**) Inactivation of *ift140* led to the formation of pronephric cysts. The glomerular neck region of the pronephros (dashed box) is dilated in the mutants. Images of embryos at 2 dpf (C) and 4 dpf (D) are shown. (**E**) Inactivation of *ift140* caused mesonephric tubular dilation. Whole-mount immunostaining was performed on optically cleared juvenile fish using an anti-PKC antibody. Open triangle, arrow, blue arrowhead, and white arrowhead indicate the glomerulus, mesonephric tubular dilation, mesonephric distal tubule, and pronephric tubule, respectively. Representative images are shown. Scale bar: 50 µm.

To assess the mesonephros, which begins to form around 12-14 dpf, we conducted optical clearing-whole mount imaging of surviving *ift140^e2/e2^* larvae fish. At 26 dpf, the mutants exhibited only a few mesonephros, compared to more than 20 in age-matched siblings, indicating a delay in the mesonephric kidney development (Figure 1E). In contrast, pronephric kidney development appeared unaffected, as demonstrated by the normal expression patterns of tubular segment markers (Supplemental Figure 3). Notably, *nephrin* expression in glomerular podocytes was expanded (Supplemental Figure 3), which aligns with the dilation observed in the glomerular-neck region (Figure 1C, 1D). The expression of *slc13a1* was also expanded, suggesting proximal tubular dilation (Supplemental Figure 3). Importantly, the mutants exhibited dilation in the newly formed mesonephric tubules (arrow), notably in the proximal region adjacent to the glomerulus but not in the distal region fused to the pronephros (blue arrow) at 26 dpf (Figure 1E). By 30 dpf, the kidneys were crowded with dilated tubules, making it difficult to determine whether the distal tubules were also dilated (Figure 1E).

Given that *IFT140* mutations are linked to several skeletal ciliopathies^26,27^, we examined craniofacial and axial skeletal development. At 6 dpf, craniofacial cartilage structures, visualized through Alcian Blue staining, were present and showed no significant morphological differences between wild-type and mutant embryos (Figure 2A). Calcified bone structures, observed via Alizarin Red staining, also appeared normal (Figure 2B). By 26 dpf, caudal non-mineralized fins, including the dorsal and anal fins, were barely detectable (Figure 2C1). Although the craniofacial cartilage structures, such as the Meckel’s (mk) and ceratohyal (ch) cartilages, were present in the mutants, they were significantly shorter compared to wild-type larvae, which might be attributed to overall small body size (Figure 2C2). In the caudal fin complex, only the hypural (hy) cartilage structure was visible, with no other cartilage structures evident (Figure 2C3). Interestingly, mutant larvae exhibited a characteristic bend in the anterior portion of the caudal fin complex (Figure 2C3). During this developmental stage, the ossified bone structures displayed the most pronounced defects in the mutants (Figure 2D1). Most craniofacial bones were absent, the number of vertebrae was greatly reduced with only a few present in the rostral part of the axial skeleton (Figure 2D2), and no structures of the caudal fin were found (Figure 2D3). While the inactivation of *ift140* may primarily contribute to these defects, developmental delay and/or global growth arrest may also play a role. Overall, our data demonstrated that zebrafish *ift140* mutants largely recapitulate the renal cystogenesis and skeletal development phenotypes observed in mammals.

**Figure 2.**
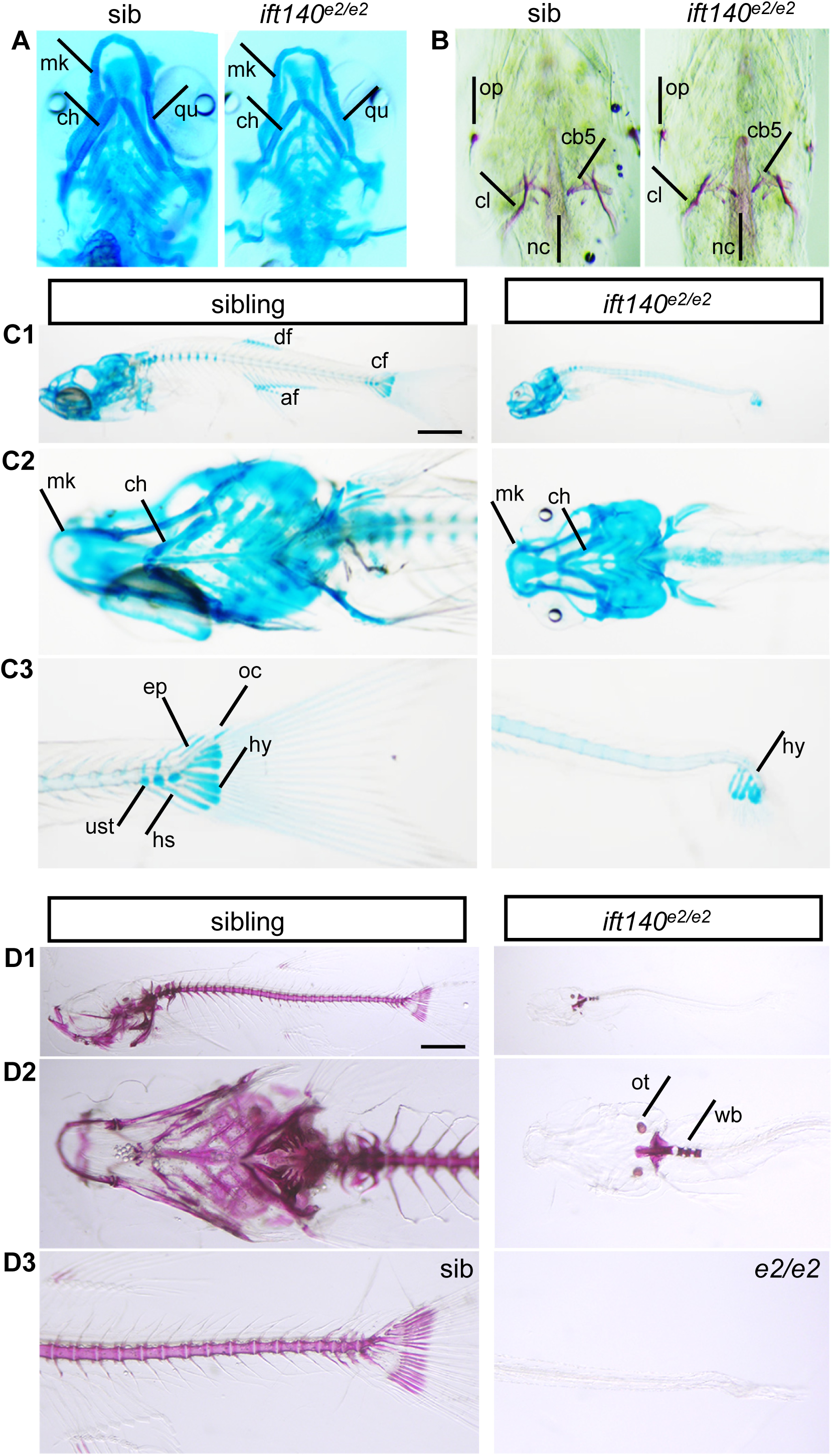
Inactivation of *ift140* results in a lack of ossification of larval bones. (**A**) Alcian Blue staining of embryos at 6 dpf shows no differences in craniofacial cartilage structures between wt and mutant zebrafish. (**B**) Alizarin Red staining of embryos at 6 dpf indicates no differences in craniofacial calcified bone structures between wt and mutants. (**C1-C3**) Alcian Blue staining of larval fish at 26 dpf reveals the presence of craniofacial cartilage structures, while caudal fin cartilage structures are absent, except for the hypural (hy). Shown are lateral views of whole larvae (C1), ventral views of craniofacial cartilage (C2), and lateral views of the caudal fin complex (C3). (**D1-D3**) Alizarin Red staining of larval fish at 26 dpf shows that most calcified bones are absent, with only a few vertebrae present in the rostral part of the axial skeleton. Shown are lateral views of whole larvae (D1), ventral views of craniofacial bones (D2), and lateral views of the caudal fin (D3). Label abbreviations: mk: Meckel’s cartilage; ch: ceratohyal; qu: palatoquadrate; op: opercle; cb5: ceratobranchial 5; nc: notochord; df: dorsal fin; af: anal fin; cf: caudal fin; hy: hypural; ep: epurals; oc: opisthural cartilage; ust: urostyle; hs: haemal spines; wb: Weberian apparatus; ot: otolith. Scale bar: 1 mm.

### *ift140^MJ^* fish enables the analysis of renal phenotypes in adult zebrafish in the F0 generation

The early death of *ift140* homozygous mutants prevented the examination of renal phenotypes in adult fish, while heterozygous *ift140* fish remained generally normal for at least one year (data not shown). To overcome these limitations, we employed F0-based mosaic knockout using MMEJ-mediated genome editing technology. The sgRNA-injected F0 fish, designated as *ift140^MJ^*, were viable with a knockout efficiency of approximately 80%, with many exhibiting significant shortening of body length and a distorted tail (Figure 3A). The kidneys of the F0 fish appeared overly wide and lost their saddle-like shape (Figure 3B), with the kidney area significantly enlarged when normalized to body length (Figure 3C). To further explore the correlation between knockout efficiency and disease severity, we titrated the sgRNA dose. We observed a dose-dependent relationship between *ift140* knockout efficiency, survival, and renal cystic burden. At approximately 50% knockout efficiency, the fish appeared largely normal, displaying only minor kidney tubular dilation. At 60-80% knockout efficiency, the fish were notably smaller and exhibited moderate kidney cysts. At knockout efficiencies exceeding 90%, most fish either died or were markedly undersized, with the kidneys filled with large numbers of cysts (Figure 3D). A Pearson’s correlation coefficient analysis between *ift140* knockout efficiencies and the kidney cystic burden revealed a strong positive relationship (r = 0.83, *P* < 0.0001), indicating a highly significant correlation (Figure 3E).

**Figure 3.**
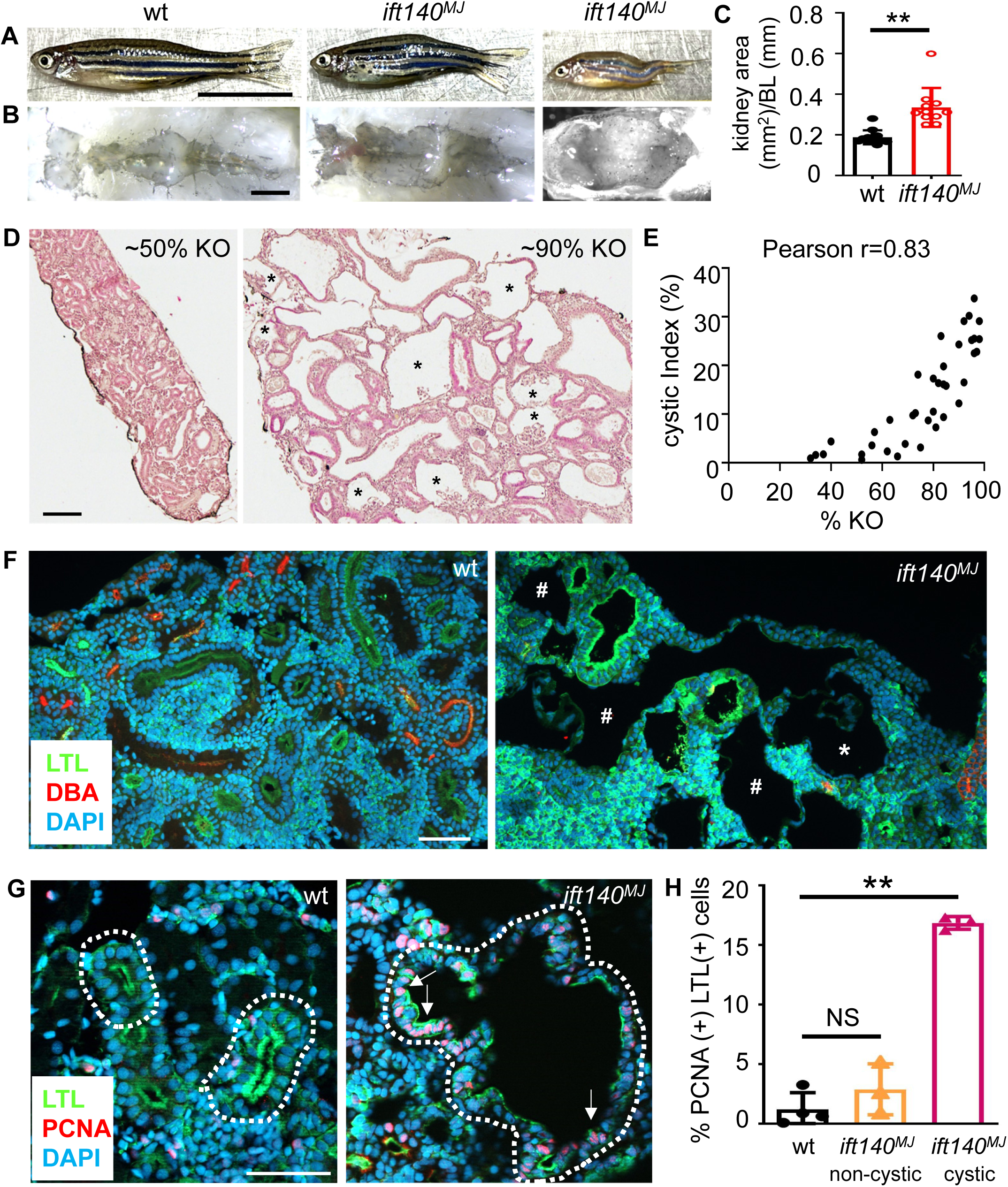
MMEJ-mediated mosaic knockout of *ift140* enables the analysis of renal cysts in adult fish in the F0 generation. (**A**) Gross morphology of adult *ift140^MJ^* fish. Embryos injected with MMEJ-inducing sgRNA targeting *ift140* were allowed to grow to adulthood. At 3 months of age, *ift140^MJ^* fish were generally shorter in length and often exhibited a distorted body shape compared with wild type fish. (**B, C**) The kidneys in adult *ift140^MJ^* fish lost their saddle-like shape (B) and became enlarged (C). The kidney area was normalized to body length (BL). Eight wild type fish and eight *ift140^MJ^* fish with a knockout efficiency (% KO) ∼80% were analyzed. % KO was calculated using genomic DNA from the tail fin. (**D, E**) Renal cyst severity correlates with % KO of *ift140*. Representative hematoxylin and eosin (H&E) images are shown, with glomerular cysts indicated by asterisks (D, *). A significant correlation between *ift140* knockout efficiencies and cyst severity was demonstrated by Pearson’s correlation coefficient (r = 0.83, *P* < 0.0001) (E). The cyst area/total kidney area (cystic index) was analyzed using three sections per kidney. (**F**) The segment identity of cystic tubules was examined by co-staining renal sections with LTL (green; labels PTs and major CDs), DBA (red; labels DTs), and DAPI (blue; labels nuclei). Unstained tubules are indicated by # , and the glomerulus is marked with asterisk (*). (**G, H**) Cyst-lining epithelial cells exhibited hyper-proliferation. Kidney sections were immunostained using a PCNA antibody (G), and the percentage of PCNA(+) LTL-labeled cells (arrows) was quantified (H). Four fish per group and three sections per kidney were used for analysis (F-H), with approximately 200 LTL-labeled cells assessed per section (H). Scale bars: 1 cm (A), 1 mm (B),100 µm (D), and 50 µm (F, G). **: *P* < 0.01; NS: not statistically significant (*P* > 0.05).

Notably, the renal cysts in the F0 *ift140^MJ^* fish included both glomerular cysts and tubular cysts (Figure 3D). To determine the origin of the tubular cysts, we stained kidney sections with LTL, which labels both proximal tubules (PTs) and major collecting ducts (CDs) in zebrafish^34^, and with DBA, which labels distal tubules (DTs) in zebrafish^34^. We found that large tubular cysts were mostly unstained by either LTL or DBA, while some medium-sized cysts were LTL (+) and no DBA (+) cysts were observed (Figure 3F). The unstained cysts are presumed to contain dedifferentiated epithelial cells. Therefore, tubular cysts originated from the PTs and possibly the CDs, where the epithelial cells became dedifferentiated following cyst expansion.

Hyper-proliferation of cyst-lining epithelial cells is a hallmark of ADPKD^3,35^. Consistently, LTL(+) cystic tubules in the *ift140^MJ^* fish exhibited significantly more PCNA-positive epithelial cells compared to those in wild-type fish (Figure 3G, 3H). Additionally, LTL(+) non-cystic tubules in the F0 animals also showed a tendency for increased cell proliferation (Figure 3H). Renal fibrosis, another hallmark of ADPKD, has been observed in *Ift140*-depleted kidneys in mice^28,36^. In line with this observation, the *ift140^MJ^* kidneys were found to be fibrotic, as evidenced by increased collagen deposition (Supplemental Figure 4).

ADPKD is a systemic disease, with cardiovascular dysfunction being a common presentation and a major cause of mortality^37,38^. Dilated cardiomyopathy has been detected in patients harboring *IFT140* pathogenic variants^39^, and laterality defects have been observed in *Ift140* mutant mice^40^. We did not observe aberrant cardiac asymmetry in *ift140* mutant embryos (data not shown). Consistent with this, the Kupffer’s vesicle (KV) cilia appeared normal (Supplemental Figure 5), although the potential influence of maternal *ift140* expression cannot be excluded. No compromised cardiac function was observed in *ift140* mutants and CRISPR-edited embryos (Supplemental Figure 6A and data not shown). However, a decline in cardiac function- was detected in adult *ift140^MJ^*fish (Supplemental Figure 6B). Together, these data illustrate the feasibility of conducting rapid genetic analyses of homozygous lethal ADPKD causative genes through MMEJ-mediated mosaic knockout technology in the F0 generation of adult zebrafish.

### Inactivation of *ift140* results in loss of cilia and aberrant epithelial polarity

To elucidate the mechanisms underlying renal cystogenesis, we examined ciliogenesis. In zebrafish embryos, the distal late pronephric tubule contains single cilium, while the distal early and proximal straight tubules are occupied by multiple cilia bundles with individual single cilium dispersed among them. The proximal convoluted tubule predominantly has single cilium, along with some cilia bundles (Figure 4A)^41,42^. In *ift140^e2/e2^* embryos, the distal single cilium was marginally shorter, and multi-cilia bundles lacked the well-aligned orientations typical of wild type cilia at 2 dpf (Supplemental Figure 7A, 7B). These defects became more pronounced by 4 dpf, with single cilia becoming even fewer in number (Figure 4B, 4B1). Furthermore, in the PST segments of mutant embryos, acetylated α-tubulin staining was noted around the cell periphery and within the cytoplasm of some cells (Figure 4D), suggesting a stabilization of intracellular microtubules.

**Figure 4.**
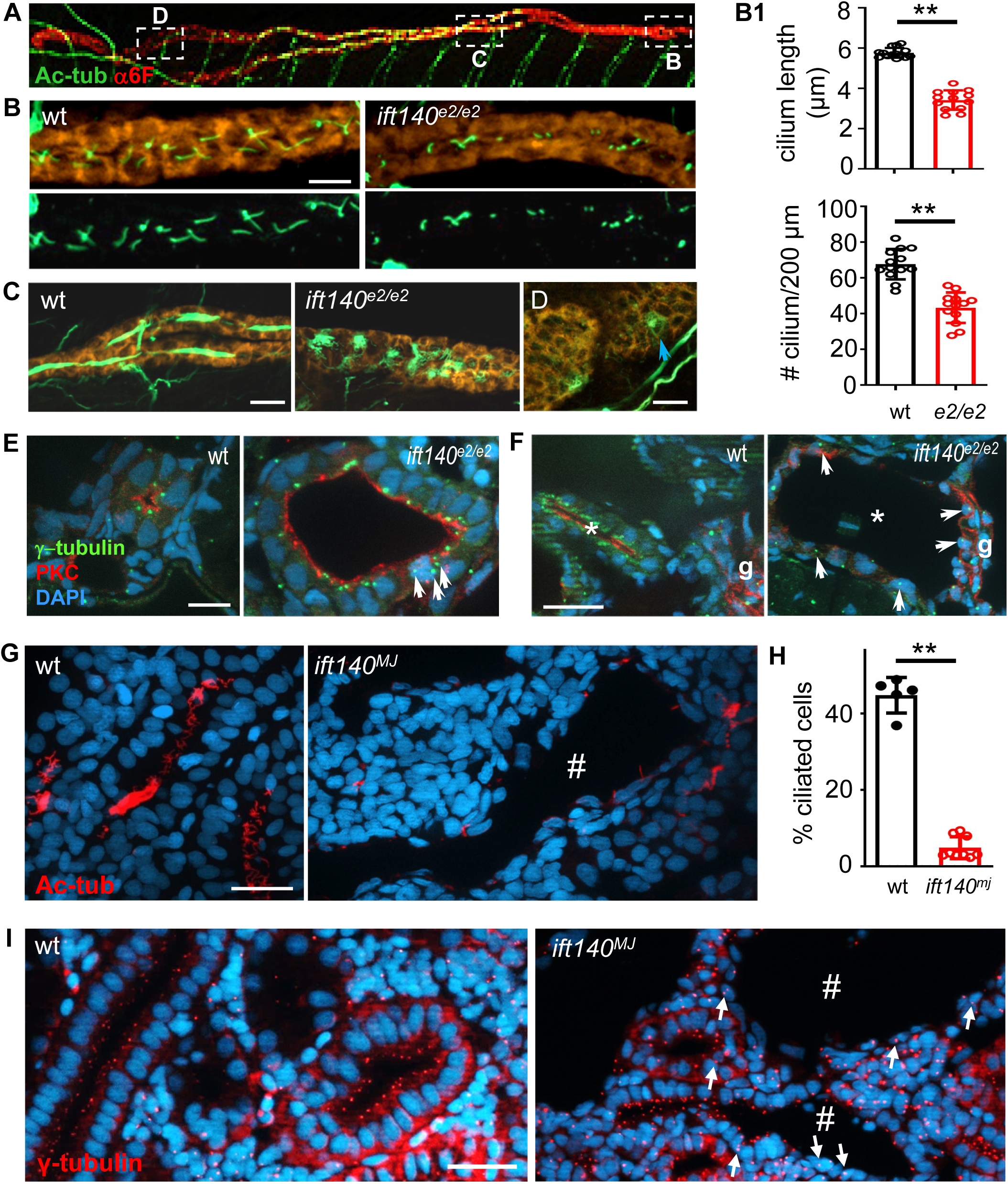
The impact of *ift140* inhibition on kidney cilia and epithelial polarity. (**A**) Illustration depicting the expression patterns of single cilia and multi-cilia bundles in a 4 dpf wt fish. Cilia were visualized by whole-mount immunostaining with an antibody against acetylated α-tubulin (Ac-tub, green), and pronephros were labeled using an antibody against Na^+^/K^+^ ATPase (α6F, red). Boxed regions are enlarged in panels (B-D). (**B, B1**) *ift140^e2/e2^* embryos displayed shorter and fewer distal single cilia. Representative images are shown in (B), with quantifications of cilium length and number provided in (B1). The total numbers of single cilia in the distal region of both pronephros were counted and normalized to the length of measured area. (**C**) *ift140^e2/e2^*embryos exhibited misoriented multi-cilia bundles. (**D**) *ift140^e2/e2^* embryos showed Ac-tub staining around the cell periphery and within the cytoplasm (blue arrow). (**E, F**) Basal body localization in *ift140^e2/e2^* embryos. Cryosections of 4 dpf embryos were immunostained using antibodies against atypical PKC (red) and γ-tubulin (green). Representative cross-section images from distal tubules (E) and the glomerular-neck region (F, asterisk) are shown. g: glomerulus. (**G-I**) Three-month-old *ift140^MJ^* kidneys exhibited fewer ciliated cells and abnormal basal body docking. Representative images of cilia staining with Ac-tub (red) are shown in (G), along with quantification of ciliated tubular epithelial cells (H). Basal body localization in tubular epithelial cells was visualized using an antibody against γ-tubulin (red) (I). The arrow indicates displaced basal body, and # marks cystic tubules. Ten fish per genotype were analyzed in panels (B-D), and 5 to 10 fish with 3 to 12 sections each were analyzed in panels (E-I), with approximately 1000 tubular epithelial cells counted (H). Scale bars: 10 µm (B, E), and 20 µm (C, D, F, G, I). **: *P* < 0.01.

The disorganization of multi-cilia bundles resembles that seen in planar cell polarity (PCP) mutants^43^. We then examined basal body docking, a process that is tightly regulated by the PCP pathway. In polarized kidney epithelial cells, the basal body is typically localized at the apical side of the cell. In *ift140^e2/e2^* embryos, the basal bodies mostly remained at the apical surface of the dilated distal tubules, with some displacement observed at the lateral junction (Figure 4E). In the severely dilated glomerular-neck region, the positions of the basal bodies became randomized, localizing at the lateral junctions, the center of the cells, or the basal surface (Figure 4F). Of note, apico-basolateral polarity, which is frequently disrupted in cystic kidney diseases, appeared unaffected in the mutant embryos, as indicated by the apical expression of atypical PKC and the basolateral localization of Na^+^/K^+^-ATPase (Supplemental Figure 8). Next, we examined ciliogenesis and basal body docking in the kidneys of adult *ift140^MJ^* fish. The single cilium was drastically depleted in the *ift140^MJ^* kidneys (Figure 4G, 4H). While multi-cilia bundles were abundant in *ift140^e2/e2^* embryos, they were significantly reduced in the adult kidneys (Figure 4G), indicating a progressive loss of cilia. Consistent with the observation in *ift140^e2/e2^*embryos, the basal bodies were largely localized to the apical side of non-cystic tubular epithelial cells but displayed randomization in cystic tubules (Figure 4I). Together, our data suggest that inactivation of *ift140* is associated with a loss of cilia, a disruption of apical docking of the basal body, and aberrant stabilization of intracellular microtubules.

### *ift140^MJ^* embryos can serve as a platform for screening protective genetic modifiers of renal cysts

Because *ift140^e2/e2^* embryos develop pronephric cysts without apparent complications, and the cyst detection method is straightforward under a light microscope from lateral and/or dorsal views at 4-6 dpf (Figure 1D, Supplemental Figure 2A), this raises the possibility that *ift140* embryos could serve as a platform for cyst-based screening of genetic modifiers of PKD. Furthermore, the injection of MMEJ-inducing sgRNA targeting *ift140* successfully recapitulated the pronephric cyst phenotype in approximately 90% of injected embryos when 5 µM of *ift140* sgRNA was utilized, resulting in knockout efficiency exceeding 90% (Figure 5A). This suggests that higher screening throughput can be achieved using *ift140^MJ^* embryos.

**Figure 5.**
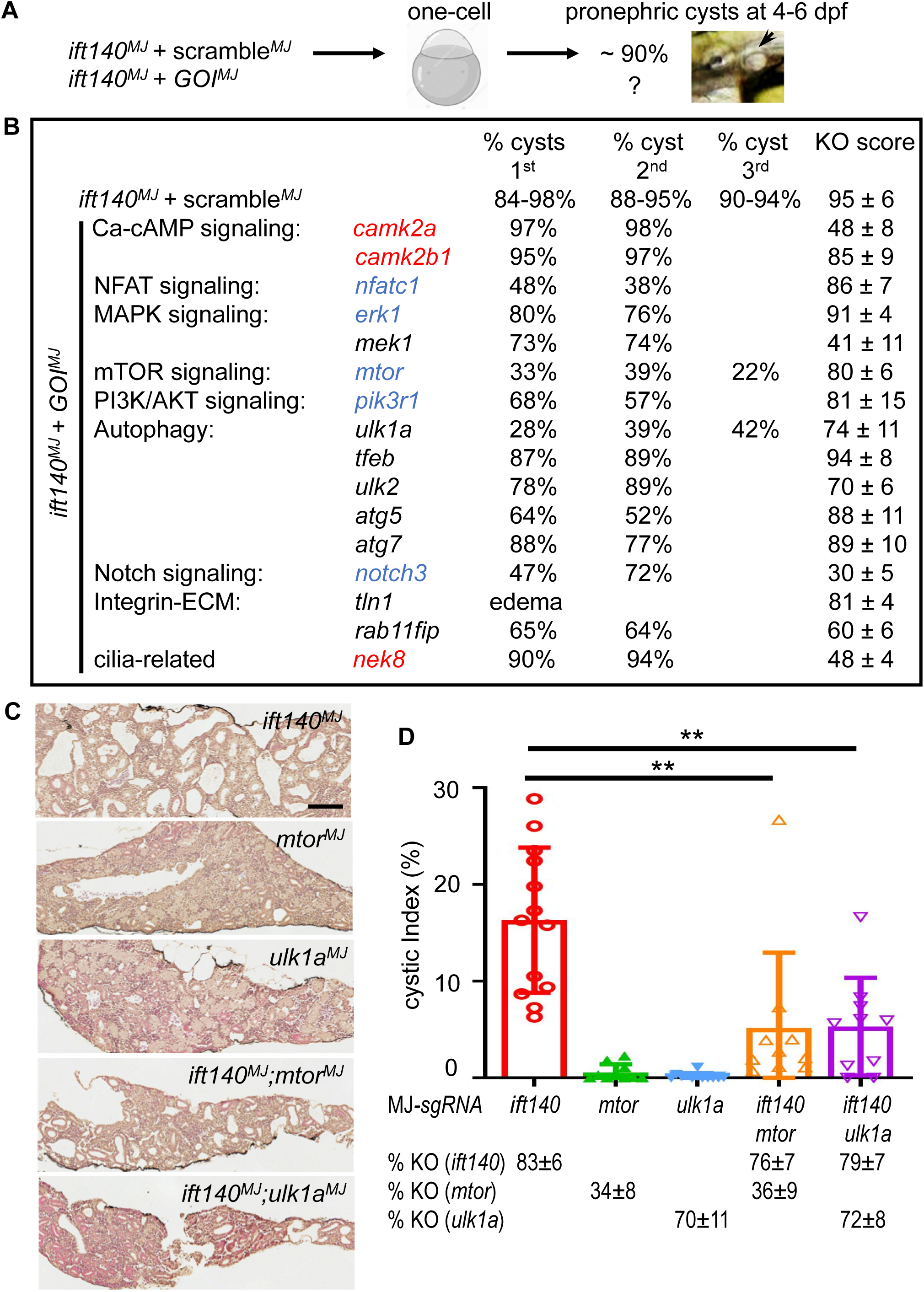
MMEJ-based F0 assay reveals protective modifying effects of known and novel genes in kidney cysts. (**A**) Schematic representation of the study design. (**B**) The effect of knocking down 16 genes in *ift140*-associated pronephric cyst formation. Each of MMEJ-inducing sgRNA targeting 16 genes of interest (GOI), implicated in dysregulated signaling pathways during cystogenesis, was co-injected with an sgRNA against *ift140* into 1-cell stage embryos. For each gene, 50-100 injected embryos were scored for pronephric cysts at 5 dpf. Injections were conducted in groups, with each group consisting of approximately 4 GOIs (*ift140^MJ^ + GOI^MJ^*) and one control (*ift140^MJ^ +* scramble*^MJ^*). After repeating experiments, gDNA was extracted from 4 embryos per gene to assess knockout efficiency. (**C, D**) Protective modifying effects of *mtor* and *ulk1a* persist into F0 adult fish. sgRNA-injected embryos were allowed to grow to 3 months of age, and their kidneys were collected for cyst analysis. Representative H&E-stained kidney images are shown in (C), with quantifications of cyst burden and knockout efficiency presented in (D). Knockout efficiency was calculated using gDNA extracted from tail fins, with a lower knockout score observed for *mtor* in surviving adult fish. Ten to twelve fish per genotype were analyzed in panel (D). Scale bar: 200 µm. **: *P* < 0.01.

We conducted a pilot screen using an MMEJ-mediated genetic assay, selecting 18 genes implicated in signaling pathways known to be dysregulated in ADPKD (Figure 5B)^44,45^. MMEJ-inducing sgRNAs targeting each of the 16 genes were co-injected with *ift140*-sgRNA into one-cell-stage embryos, and the pronephric cysts in the co-injected embryos were assessed at 4-6 dpf and compared with those injected with *ift140*-sgRNA and a scramble sgRNA (Figure 5A, 5B). Indeed, inactivation of the genes highlighted in blue, which were previously suggested to have beneficial effects based on genetic or pharmacological studies, reduced the number of *ift140^MJ^* embryos with pronephric cysts, although some effects were marginal^46–48^. In contrast, inactivation of the genes marked in red, which were previously suggested to be detrimental, did not suppress pronephric cyst formation in *ift140^MJ^* embryos^49–52^. Importantly, in addition to genes with known modifying effect such as *mtor*, the pilot screen also suggested novel candidate protective modifiers, such as *ulk1a* (Figure 5B). We repeated the injections and obtained largely reproducible results (Figure 5B).

To determine whether the modifying effects of *mtor* and *ulk1a* persisted into adulthood, we allowed the injected embryos to grow. While the majority of *ift140^MJ^;mtor^MJ^*and *ift140^MJ^;ulk1a^MJ^* fish died, the survivors exhibited significantly less cyst burden than *ift140^MJ^* fish at 3 months (Figure 5C, 5D). Of note, samples with similar *ift140* knockout efficiency were selected for comparison to ensure that the observed modifying effects were not due to varying degrees of *ift140* knockdown (Figure 5D)

To confirm the modifying effects of *mtor* and *ulk1a* observed in the F0-based assay, we utilized stable mutants for *mtor* and *ulk1a* that were generated previously^25,53^. First, *ift140*-sgRNA was injected into embryos obtained from crosses between *mtor^+/-^* and wt fish. At 5 dpf, *ift140^MJ^*;*mtor^+/-^* embryos developed significantly fewer pronephric cysts than their sibling *ift140^MJ^* fish (Figure 6A). Similarly, *ift140*-sgRNA was injected into embryos obtained from crosses between *ulk1a^+/-^* and wt fish, and the percentage of *ift140^MJ^;ulk1a^+/-^*embryos exhibiting pronephric cysts was significantly reduced compared to that of their sibling *ift140^MJ^* embryos (Figure 6B). Next, we examined the *ift140*;*mtor* and *ift140*;*ulk1a* stable double mutants. Unlike the *ift140^MJ^;mtor^+/-^* fish, *ift140^-/-^;mtor^+/-^* fish still formed cysts (Figure 6C), but these cysts appeared smaller (data not shown), which could be attributed to differences in knockout efficiency between *ift140^MJ^* and *ift140^-/-^*. However, *ift140^-/-^;mtor^-/-^* embryos showed a significant reduction in cyst formation (Figure 6C). On the other hand, both *ift140^-/-^;ulk1a^+/-^*and *ift140^-/-^;ulk1a^-/-^* embryos were protected from cyst development (Figure 6D). We thus conclude that protective modifiers for *ift140*-based kidney cysts can be effectively identified using the F0-based genetic assay and can be confirmed in stable mutants.

**Figure 6.**
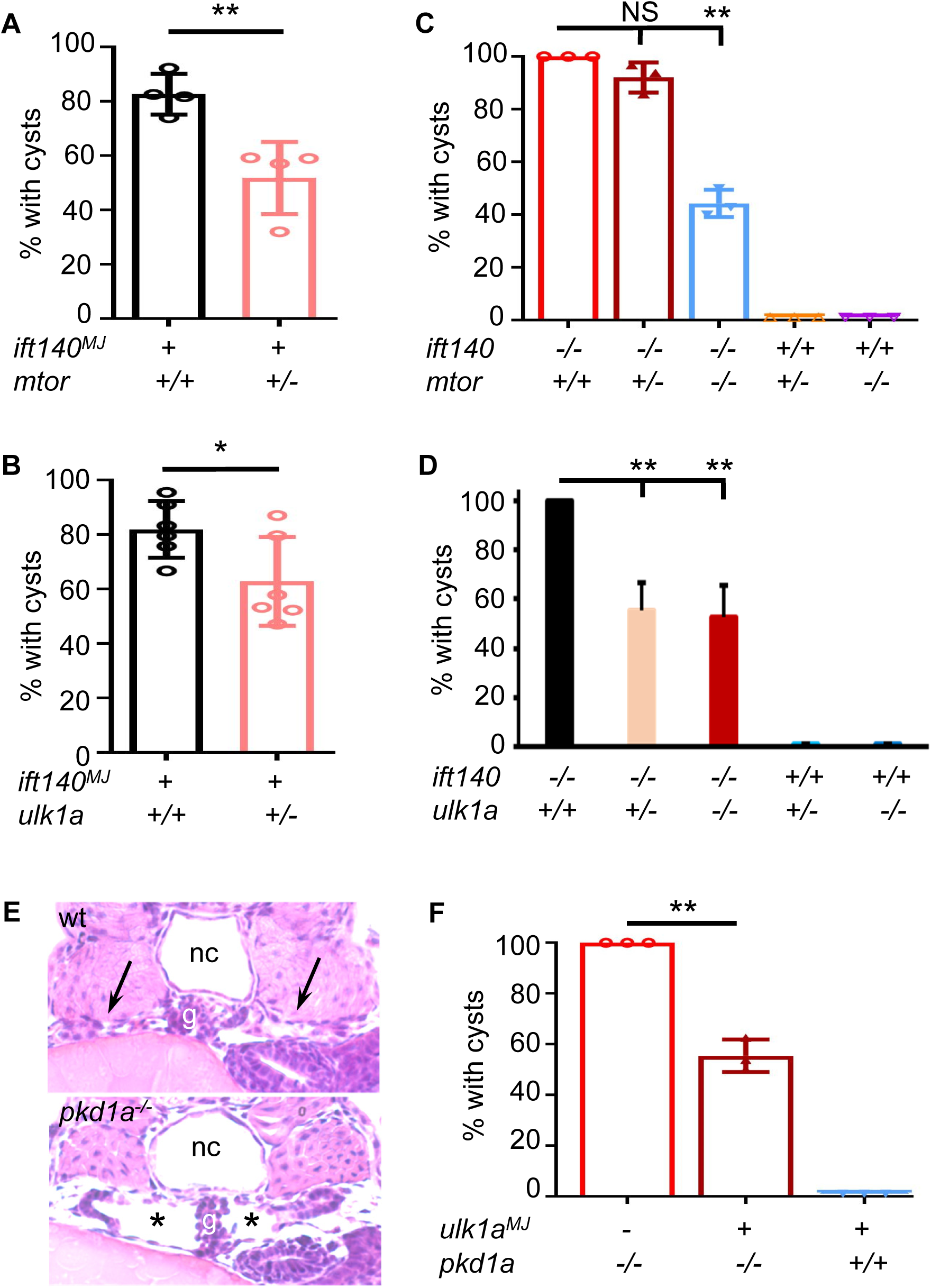
The protective role of *mtor* and *ulk1a* inhibition is validated via stable mutants and extends to *pkd1^-/-^* embryos. (**A, B**) Haploinsufficiency of *mtor* or *ulk1a* protects against *ift140^MJ^* - associated pronephric cyst development. sgRNA targeting *ift140* was injected into embryos obtained from crosses between *mtor^+/-^* and wild-type fish or between *ulk1a^+/-^* and wild-type fish. At 5 dpf, pronephric cysts were scored, and all embryos were genotyped for *mtor* (A) or *ulk1a* (B). (**C, D**) Haploinsufficiency of *mtor* or *ulk1a* protects against *ift140^e2/e2^*-associated pronephric cyst formation. Embryos obtained from *ift140^+/-^;mtor^+/-^*inter-crosses or *ift140^+/-^;ulk1a^+/-^* inter-crosses were scored for pronephric cysts at 5 dpf and genotyped for *ift140* and *mtor* (C) or *ift140* and *ulk1a* (D). (**E, F**) Knockdown of *ulk1a* protects against *pkd1^-/-^* - associated pronephric cyst formation. sgRNA targeting *ulk1a* was injected into embryos from *pkd1^+/-^* inter-crosses. At 3 dpf, embryos were fixed in 4% PFA, and posterior tails were collected for genotyping. Cyst formation was assessed by H&E staining of JB- 4 sections. Representative images show the glomerulus-neck region of the pronephros in wild-type embryos (arrowhead) and *pkd1^-/-^* embryos (asterisks indicate dilation). Results are from three independent experiments, with > 100 embryos analyzed per experiment (A, B) and 7–20 (C, D) or 6–10 (F) embryos per genotype examined per experiment. *: *P*<0.05. **: *P*<0.01. NS: not statistically significant (*P*>0.05). g: glomerulus. nc: notochord.

To investigate whether *ift140*-based modifiers are applicable to *pkd1*-based models, we injected the sgRNA targeting *ulk1a* into *pkd1^-/-^* embryos and found that inactivation of *ulk1a* was protective (Figures 6E, 6F). In conjunction with our earlier observation that inhibition of *mtor* prevents cyst formation in *pkd1^-/-^* embryos^14^, our data suggest that at least some modifier genes identified from the *ift140*-based screen could also be effective in *pkd1*-based cystogenesis.

### Inhibition of *mtor* and *ulk1a* both restores ciliogenesis and cilia orientation

To decipher the mechanisms underlying the protective effects of *mtor* and *ulk1a* inhibition, we examined ciliogenesis and cilia orientation. Consistent with the observation in *ift140^e2/e2^* mutants, *ift140^MJ^* embryos exhibited reductions in the length and number of distal single cilia (Figure 7A, 7B), misorientation of multi-cilia bundles (Figure 7C), and accumulation of acetylated tubulins within the cells (Figure 7D). However, injection of sgRNAs targeting either *mtor* or *ulk1a* rescued all of these abnormalities in *ift140^MJ^* embryos, while knockdown of *mtor* or *ulk1a* alone had no apparent effects (Figure 7A- 7D). These data suggest that *mtor* and *ulk1a* perform similar functions in the regulation of cilia size, cilia orientation, and microtubular stability.

**Figure 7.**
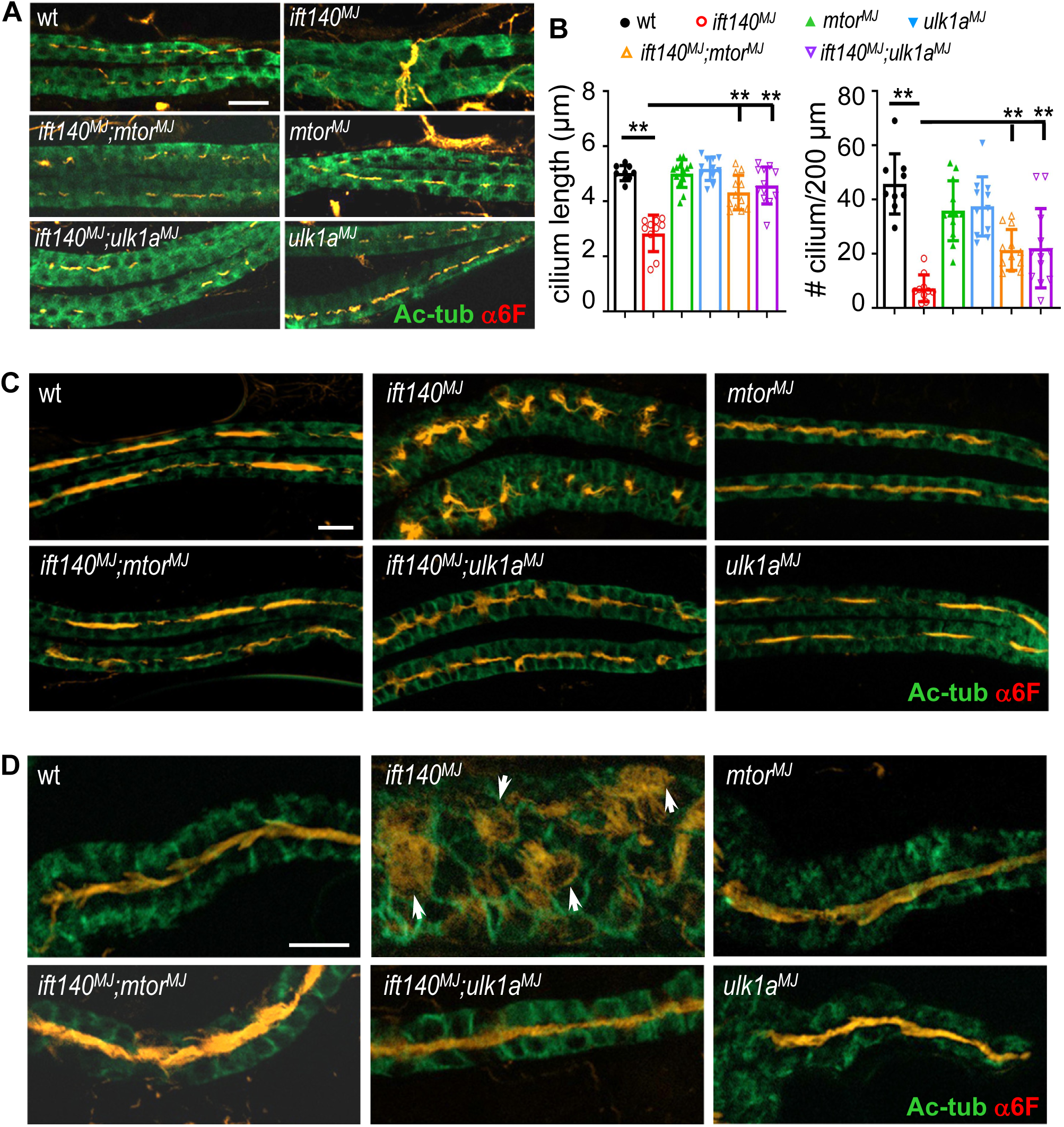
Inhibition of *mtor* and *ulk1a* restores ciliogenesis and corrects misorientation of multi-cilia bundles in *ift140^MJ^*embryos. (**A, B**) Knockdown of *mtor* or *ulk1a* both rescued the length and number of distal single cilia in *ift140^MJ^*embryos. sgRNAs targeting *ift140*, *mtor*, or *ulk1a* were injected into embryos either individually or in combination, and distal tubular single cilia were analyzed. Representative images of Ac-tub and α6F immunostaining are shown in (A), with quantifications of cilium length and number presented in (B). (**C**) Knockdown of *mtor* or *ulk1a* normalized the orientation of multi-cilia bundles in the distal tubules of *ift140^MJ^*embryos. (**D**) Knockdown of *mtor* or *ulk1a* eliminated aberrant Ac-tub accumulation (arrow) within the proximal tubule cells of *ift140^MJ^*embryos. A total of 8-12 embryos per group at 6 dpf were analyzed across two independent experiments. Scale bar: 20 µm. *: *P* < 0.05. **: *P* < 0.01.

Interestingly, *mtor* and *ulk1a* had differential effects on body length regulation in the context of *ift140* deficiency. In the same clutches of injected embryos used for cilia analysis (Figure 7), *ift140^MJ^*larvae were significantly smaller than un-injected controls at 2 weeks of age (Supplemental Figure 9A). Knockdown of *ulk1a* promoted body growth in *ift140^MJ^* fish, whereas knockdown of *mtor* did not yield the same effect, as *mtor* knockdown alone suppressed fish growth (Supplemental Figure 9A). Corresponding with the body length phenotype, the ossification of bone structures was repaired by *ulk1a* inactivation but not by *mtor* inactivation (Supplemental Figure 9B).

## Discussion

### Zebrafish *ift140* mutants and CRISPRants recapitulate the major phenotypes observed in mammalian models

Recently, *IFT140* is identified as a causative gene for ADPKD, particularly enriched in patients with milder disease presentations and no known family history of ADPKD^30,54–57^. In the mice, homozygous missense mutation of *Ift140* results in embryonic lethality. By the time of death at approximately E13.5, which is prior to the earliest cyst formation at E15.5 in *Pkd1^-/-^* embryos, the kidneys in the mutants were generally comparable to those of wild-type siblings^29,58^. However, kidney collecting duct-specific knockout of *Ift140* resulted in rapidly progressive cystogenesis^28^. In zebrafish, homozygous *ift140* mutants exhibit pronephric and mesonephric cysts but are juvenile lethal. In contrast, *ift140* CRISPants effectively overcome the early lethality, allowing for the assessment of *ift140* function in adult zebrafish. Renal cysts were clearly derived from PTs but not DTs. Determining whether cysts also originated from CDs will require the identification of a zebrafish CD-specific marker in future studies. Interestingly, adult *ift140* CRISPRants exhibited significant numbers of glomerular cysts. Though the presence of glomerular cysts in *IFT140-*ADPKD patients remains to be investigated, this phenotype is commonly observed in early-onset ADPKD cases and have also been reported in adult ADPKD, including mouse models and patients^59–62^. The zebrafish *ift140* model provides a valuable platform for further mechanistic studies.

*IFT140* was originally identified as a causative gene for skeletal ciliopathy under recessive conditions, such as SRTD9. The skeletal features of SRTD9 commonly include a small thoracic cage (short ribs), shortened long bones, and phalangeal cone-shaped epiphyses, with occasional short stature^26,27^. In missense mutant mice, severe craniofacial defects, such as agenesis or hypoplasia of facial bones, and severe rib abnormalities, including vertebral fusion, lateral branching and thickening, have been reported^29^. In zebrafish *ift140^e2/e2^* mutants, skeletal development appeared normal up to 7 dpf. However, from 7 dpf onward, the formation of calcified bone structures, including craniofacial bones, vertebrae, and caudal fins, was halted. Mutant fish also exhibited arrested growth, ultimately leading to premature death at the juvenile stage. Overall, zebrafish *ift140* mutants recapitulate many of the skeletal features observed in mammals with *IFT140* mutations, though the phenotypes appear more severe, likely due to the null mutant nature of the zebrafish model. Nonetheless, this model could be valuable for studying skeletal development associated with cilia signaling.

Using MMEJ-mediated mosaic knockout for *ift140*, our data demonstrate the feasibility of performing dose-dependent genetic analysis of ADPKD causative genes in the F0 generation of adult zebrafish. This approach enables the study of adult phenotypes that are otherwise missed in heterozygous mutants due to slow disease progression or in homozygous mutants due to embryonic or juvenile lethality, by fine-tuning the knockout efficiency of a gene of interest. The MMEJ method not only offers a rapid alternative to tissue-specific knockout of lethal genes but also better replicates the whole-body genetic lesions observed in human patients. For non-lethal mutations, this technology can significantly expedite experimental timelines by eliminating the need for multi-generation crosses. One limitation of this approach is the potential for unclean genetic lesions, including off-target effects of sgRNA injections. To mitigate this, using multiple sgRNAs targeting the same gene and comparing the results can help differentiate shared phenotypes from unique ones.

### Both ciliary and non-ciliary roles of IFT140 might be implicated in renal cyst formation and can be studied in zebrafish

The mechanisms underlying *IFT140*-associated cystogenesis remain unclear. IFT140 is a component of the IFT-A complex, which facilitates retrograde trafficking of cargos from the cilium tip to the base. Consistent with this role, mutations in *IFT140* result in shortened cilia and abnormal accumulation of ciliary proteins, as observed in this and other studies^27–29^. Moreover, IFT-A complex proteins are also implicated in anterograde transport of materials into the cilium. For instance, *Drosophila ift140* is required for the ciliary localization of a TRPV calcium channel^63^; *C. elegans ift140* regulates the ciliary entry of transition zone proteins^32^; and mammalian *IFT140* controls the entry of G protein-coupled receptors into cilia^64^. Altered cilia-mediated signaling, including disrupted ciliary localization of PC1 or PC2, may underlie renal cyst development. However, *ift140* zebrafish mutants exhibit fully penetrant pronephric cyst formation as early as 2 dpf, despite only marginally shortened cilia at that stage. This observation raises the question of how much cilia length, per se, contributes to cyst formation.

In addition to their well-established roles in ciliary functions, growing evidence highlights the non-ciliary roles of IFT proteins. For instance, *IFT20* and *IFT88* are essential for the proper orientation of the mitotic spindle^65–68^. The defects in spindle orientation could affect proper cell division and PCP and thus contribute to renal cyst formation. Furthermore, depletion of *IFT88* increases cytoplasmic α-tubulin acetylation, such a posttranslational modification is associated with stabilized, long-lived microtubules^69^. Mutations in *IFT54* and *IFT52* lead to similar microtubule stabilization^68,70,71^. The defects in cytoplasmic microtubule dynamics were shown to affect cytoskeletal architecture, intracellular trafficking of proteins, and polarity alteration in renal cells^70^, and were also noted in ARPKD patient samples^69^, suggesting that the non-ciliary roles of IFT proteins, particularly those within the IFT-B complex, may also contribute to the development of renal cysts.

Kidney collecting duct-specific knockout of *Ift140* leads to renal cyst formation without affecting mitotic spindle orientation or the apical localization of centrosomes^28^. In our study, we observed randomization of basal body docking in *ift140* mutants and CRISPRants, particularly in the proximal tubular segment. Additionally, increased α- tubulin acetylation was noted within the proximal tubular cells, closely resembling findings in ARPKD patients, cultured cells with *IFT88* depletion or mutations in *IFT54* and *IFT52*, as well as zebrafish *ift52* mutants^68–71^. This abnormality in microtubule dynamics may partially explain the misorientation of multi-cilia bundles and the randomization of basal body docking observed in the proximal tubules of *ift140* mutants and CRISPRants. Further studies are needed to investigate whether altered microtubule dynamics are present in human patient samples and to reconcile the apparent discrepancy in PCP-related phenotypes between our zebrafish mutants and mouse knockouts, potentially attributed to differences in tubular segment types (proximal tubule vs. collecting duct). Notably, alterations in epithelial morphology and cytoskeletal architecture have been documented in the limb bud of mouse *Ift140* missense mutants, as evidenced by the delocalization of epithelial cadherin (E-cad) and filamentous actin (F-actin)^29^. In summary, our study is the first to suggest the non-ciliary roles of an IFT-A protein in kidney cyst development, justifying the need for future studies to explore its contribution to cystogenesis.

### *ift140* CRISPRants provide a useful platform for the assessment of genetic modifiers of kidney cysts

The high penetrance and straightforward detection of *ift140*-associated cysts, combined with the high efficiency of MMEJ-mediated gene knockdown, led us to explore whether the *ift140* model could be used for cyst-based genetic screening in the F0 generation. By testing 16 genes implicated in signaling pathways known to be dysregulated in ADPKD^44,45^, our findings suggest that MMEJ-based F0 screening with *ift140* embryos could serve as a valuable platform for larger-scale genetic screens in the future.

The zebrafish embryonic cyst model offers unparalleled high throughput; however, the extent of its conservation with the pathogenesis of mammalian PKD requires further evaluation. While mTOR inhibition has been reported to provide benefits across various animal models of kidney cysts, the role of Ulk1a inhibition has not been previously explored. Future studies examining the effects of these interventions in a mouse *ift140* model, ideally a viable hypomorphic mutant with global disruption of *ift140*, will help determine the degree of conservation between the zebrafish embryonic model and mammalian PKD. Although the role of cilia motility in the embryonic kidney, which is closely associated with cyst formation, has raised concerns about the applicability of zebrafish embryos for modeling mammalian PKD, certain aspects, such as altered cilia-associated signaling in cystogenesis, may be conserved. Our findings also suggest that the non-ciliary functions of IFT140, particularly in cytoskeletal microtubule dynamics and cytoskeletal architecture, play at least some role in cyst formation or expansion. Further investigations to distinguish the ciliary and non-ciliary roles of IFT140 in renal cystogenesis could enhance the credibility of *ift140*-based embryonic screens for genetic modifier discovery.

Mutations in *PKD1* account for approximately 85% of ADPKD cases, making the discovery of modifiers for *PKD1*-associated ADPKD particularly impactful. Our findings show that the protective effects of mTOR and Ulk1a inhibition extend to *pkd1* zebrafish; however, not all modifiers identified in the *ift140*-based screen are applicable to *pkd1*- based models. For instance, knockdown of *atg5*, which has been shown to exacerbate *pkd1*-associated pronephric cysts via morpholino-directed gene disruption^14^, mitigates cyst formation in *ift140*-associated cysts. Despite these differences, the higher throughput and partial overlap of shared modifiers suggest that *ift140*-based screens could still serve as an effective platform for identifying modifiers of *pkd1*-based cystogenesis. In summary, we anticipate that the MMEJ-based F0 assay will enable the experimental evaluation of a larger pool of candidate cyst-causing genes, accelerating the study of genetic factors contributing to ADPKD.

## Methods

### Sex as a biological variable

For experiments using zebrafish embryos or larva, clutch-matched siblings were used without sex bias because it is impossible to distinguish animal sex at these stages. For experiments using young adult zebrafish, both males and females were examined, and similar results were obtained. As sex was not considered as a biological variable, pooled data from both sexes were used for analysis.

### Generation of zebrafish ift140 mutants

*ift140* genomic lesion was generated via MMEJ-mediated genome editing technology ^22,25^. F1 fish carrying germ-line mutations were identified using Conformation Sensitive Gel Electrophoresis (CSGE) on PCR products that cover the predicted indels, and confirmed by Sanger sequencing (Genewiz)^72^. Genotyping was performed using dCAP (derived cleaved amplified polymorphic sequences) primers to create a NcoI restriction enzyme site in the PCR product of mutant allele^73^. The desired F1 fish were further outcrossed to diminish the risk of off-target effects in subsequent generations. The fish used in experiments were of F3 or F4 generations. Sequences of sgRNA and primers can be found in Supplemental Table 1.

### MMEJ-based F0 assay in zebrafish embryos and adult fish

MMEJ-inducing sgRNAs for 16 genes of interest (Supplemental Table 1) were designed using the online tool MENTHU (http://genesculpt.org/menthu/) and synthesized from Synthego (Synthego Corporation) as previously described^23,24^. Each sgRNA was mixed with Alt-R Cas9 protein (Integrated DNA Technologies, 1081058) to achieve a final concentration of 5 µM and 300 ng/µL respectively, in a buffer consisting of 20 mM HEPES, 100 mM NaCl, 5 mM MgCl_2_, and 0.1 mM EDTA at pH 7.5. Approximately 3 nL of the sgRNA-Cas9 complex (sgRNP) was injected into one-cell staged zebrafish embryo, and the resulting embryos were scored for pronephric cysts at 5 dpf. In some cases, the injected embryos were allowed to grow to adulthood for the assessment of adult fish kidney phenotypes. The efficiency of the sgRNAs was assessed using genomic DNAs extracted from injected embryos or the tail fin of adult fish and calculated by the Inference of CRISPR Edits (ICE) Tool (https://www.synthego.com/products/bioinformatics/crispr-analysis). The sequences of the PCR primers utilized for evaluating KO efficiency are provided in Supplemental Table 1.

### Histological analysis

Adult zebrafish kidneys were collected as previously described^74^. Briefly, fish were euthanized using a 0.2% Tricaine solution, the internal organs were removed, leaving the kidneys attached to the dorsal abdominal wall. The fish were then fixed in 4% formaldehyde (PFA) solution overnight at 4°C. The kidney was carefully excised the following day and subsequently processed for paraffin embedding and H&E staining. Cyst burden was quantified as a percentage of cyst area over the total kidney tissue area using ImageJ software.

### Alcian Blue and Alizarin Red staining

Cartilage staining and mineralized bone staining of zebrafish larvae was adapted from published protocols^75,76^. In brief, after euthanasia with MS222 (0.02%), larvae were fixed in 4% PFA overnight at 4^0^ C. Samples were washed three times with phosphate-buffered saline with 0.1% Tween-20 (PBST) and then bleached in 1.5% H_2_O_2_ and 1% KOH for 20 minutes. For cartilage staining, samples were immersed in 0.1% Alcian Blue (w/v) (Sigma-Aldrich) in 70% ETOH overnight. For skeletal staining, specimens were incubated with 30% saturated borax for 5 hours. This was followed by incubation with 0.01% Alizarin Red (w/v) (Sigma-Aldrich) in 1% KOH overnight. The specimens from both stainings were then cleared using successive changes of 20% glycerol (Thermo Fisher) in 1% KOH, 50% glycerol in 1% KOH, and 80% glycerol in 1% KOH. Imaging was conducted within 3 days after staining using a Leika dissecting microscope with digital camera attached.

### Immunofluorescence labeling

Immunofluorescence analysis of zebrafish pronephros was performed by whole-mount immunostaining of embryos to visualize cilia, or on cryosectioned materials to visualize basal body positioning, as previously described^14,74,77^. Immunofluorescence analysis of adult zebrafish kidneys was conducted on paraffin sections as described^74^. Briefly, 20 µm- thick sections were labeled with renal tubular segment marker rhodamine DBA and LTL (Vector Laboratories). Nuclei were stained with DAPI (Vector Laboratories). Primary antibodies used include anti-acetylated α-tubulin (Sigma-Aldrich, T7451), anti-ɣ-tubulin (Sigma-Aldrich, T5326), anti-PKC (Santa Cruz Biotechnology, sc-216), anti-PCNA (Sigma-Aldrich, p8825), and anti-Na^+^K^+^ ATPase alpha-1 subunit (α6F, Developmental Studies Hybridoma Bank). Secondary antibodies are Alexa Fluor conjugated (Life Technologies). Images were captured using a Zeiss Axioplan II microscope equipped with ApoTome and AxioVision software (Carl Zeiss Microscopy)

### Optical clearing and whole-mount immunostaining of juvenile zebrafish

Juvenile zebrafish at 20-30 dpf were fixed in 4% PFA overnight and treated with a bleaching solution (1.5% H_2_O_2_ in 1% KOH) to eliminate melanocyte pigmentation. Then, whole-mount immunostaining and imaging was conducted as previously described^74^. Briefly, the fish were immersed in a hydrogel solution containing 0.25% X-CLARITY polymerization initiator overnight at 4°C, transferred to the X-CLARITY™ Polymerization System under vacuum for 3 hours at 37°C to facilitate hydrogel polymerization, and then incubated with tissue clearing solution (8 g SDS, 1.25 g boric acid in 100 ml water, pH 8.4) with shaking for 3 days (Logos Biosystems C20001). Following the clearing process, samples were washed in PBS and incubated with anti-PKC antibody and Alexa Fluor anti-rabbit IgG 568 (Invitrogen, A11011). The cleared and stained samples were mounted using the X-CLARITY Mounting Solution (Logos Biosystems, # C13101). Imaging was performed using a Zeiss LSM 780 microscope and Zen software (Carl Zeiss Microscopy).

### Western blotting

Juvenile zebrafish were homogenized in RIPA lysis buffer (Sigma-Aldrich, R0278) as described previously^14^. Antibodies used include anti-Ift140 (Proteintech®, 17460-I-AP) and anti-Actin (Sigma-Aldrich, A3854).

### Statistics

Data were presented as mean ± SD and analyzed using GraphPad Prism software. To compare differences between 2 groups, an unpaired 2-tailed Student’s *t* test was used; to compare differences among multiple groups, 1-way ANOVA followed by Tukey’s post hoc test was performed, as described in the figure legends. A *P* value of less than 0.05 was considered as statistically significant. All data collection and analysis were conducted in a blinded fashion to offset individual bias.

### Study approval

Zebrafish (*Danio rerio*) was maintained under standard laboratory conditions. All protocols and procedures were approved by the Mayo Clinic Institutional Animal Care and Use Committee (Rochester, Minnesota, USA).

### Data availability

Data values for all individual points in graphs are available in the supplemental Supporting Data Values file.

## Author contributions

PZ contributed to conducting experiments, acquiring data, analyzing data, and writing the manuscript. AL contributed to conducting experiments and acquiring data. XX contributed to study design and manuscript editing. XL contributed to study design, methodology, data analysis, and manuscript writing and editing.

## Acknowledgements

This work was supported by NIH R56 DK137778 (XX and XL), Mayo PKD Center Zell PKD Research Innovation Fund (XL), and the Mayo Foundation for Medical Education and Research (XX)

