## supplemental materials for "The *ift140*-Deficient Zebrafish: A Model for Renal Cystogenesis and an F0-Based Screen to Identify Genetic Modifiers of Kidney Cysts"

Supplemental Figure 1. *ift140* expression in zebrafish embryos
Supplemental Figure 2. *ift140*$^{e2/e2}$ fish exhibit significantly smaller size
Supplemental Figure 3. Inactivation of *ift140* does not affect kidney development
Supplemental Figure 4. Fibrosis in the kidneys of adult *ift140*$^{MJ}$ fish
Supplemental Figure 5. *ift140*$^{e2/e2}$ embryos display normal cilia in the Kupffer's vesicle
Supplemental Figure 6. Cardiac phenotypes in *ift140*$^{e2/e2}$ embryos and adult *ift140*$^{MJ}$ fish
Supplemental Figure 7. The effect of *ift140* inactivation on the length of distal single cilia and the orientation of multi-cilia bundles in the 2 dpf pronephros
Supplemental Figure 8. *ift140*$^{e2/e2}$ embryos exhibit normal apico-basolateral polarity in kidney tubular epithelial cells
Supplemental Figure 9. Effects of *ulk1a* and *mtor* inhibition on body size and calcified bone structures in *ift140*$^{MJ}$ larvae
Supplemental Table 1: List of the sequences for F0-based MMEJ screen and primers for quantifying knockout score

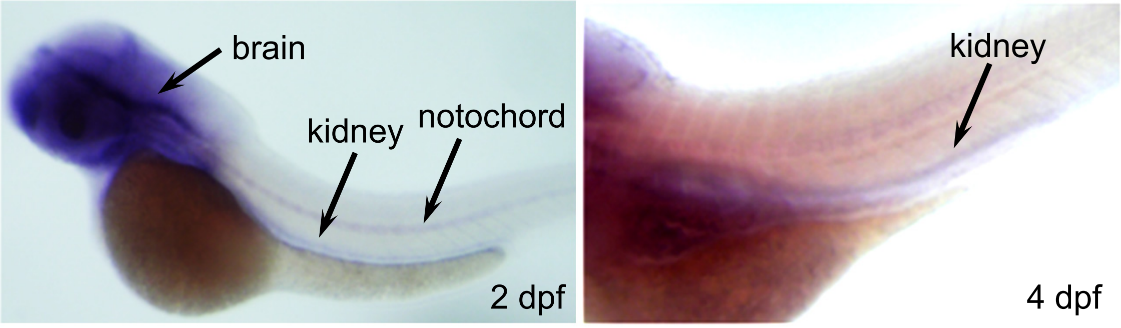


**Supplemental Figure 1. *ift140* expression in zebrafish embryos**
*In situ* hybridization reveals *ift140* transcripts localized in the pronephric kidney, notochord, and brain at 2 dpf and 4 dpf.

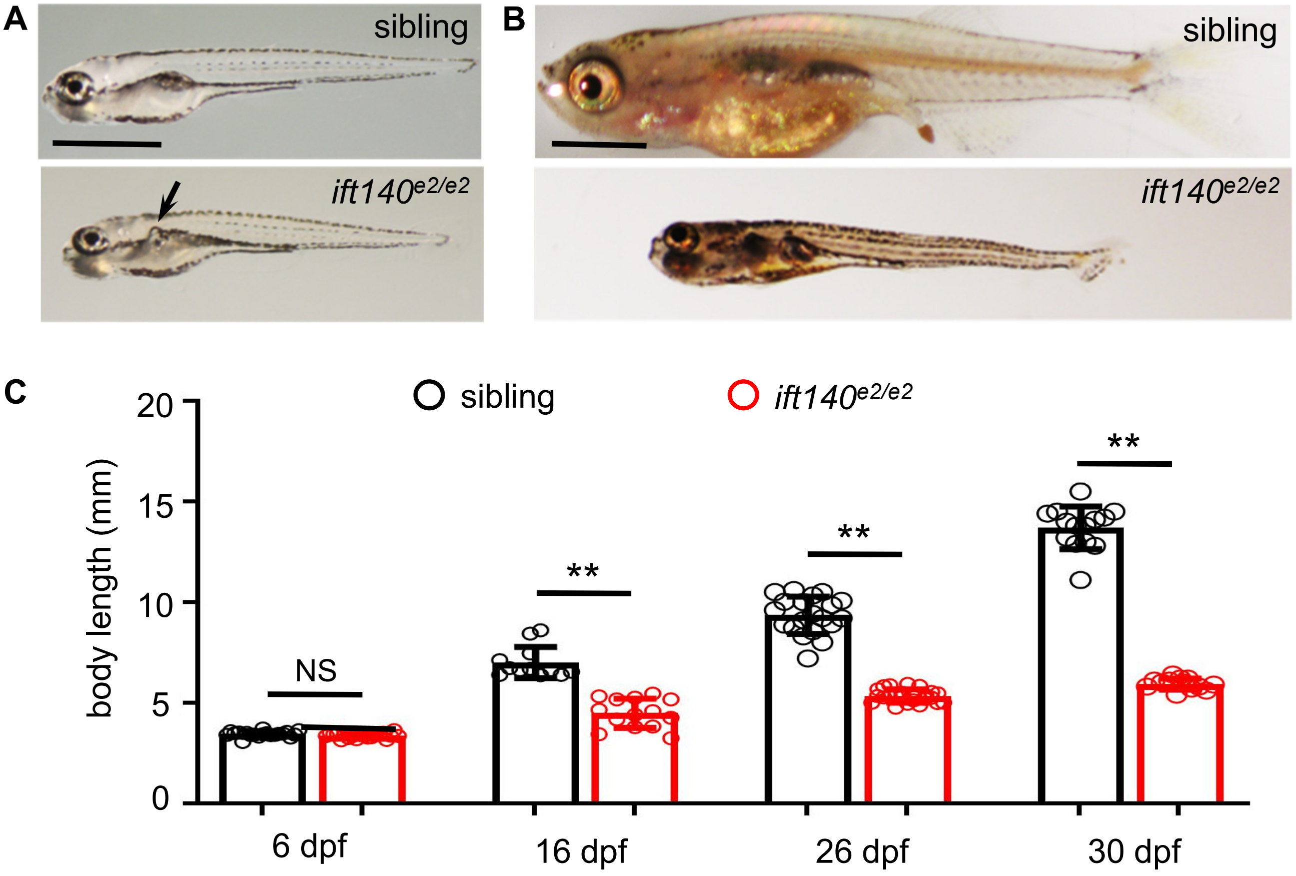


**Supplemental Figure 2. *ift140$^{e2/e2}$* fish exhibit significantly smaller size**
(**A**) *ift140$^{e2/e2}$* fish have comparable body lengths to their siblings at 6 dpf. (**B**) *ift140$^{e2/e2}$* fish are significantly smaller than their siblings from 16 dpf onward. (**C**) Quantification of body length. Scale bar: 1 mm. **: $P$<0.01. NS: not statistically significant ($P$>0.05).

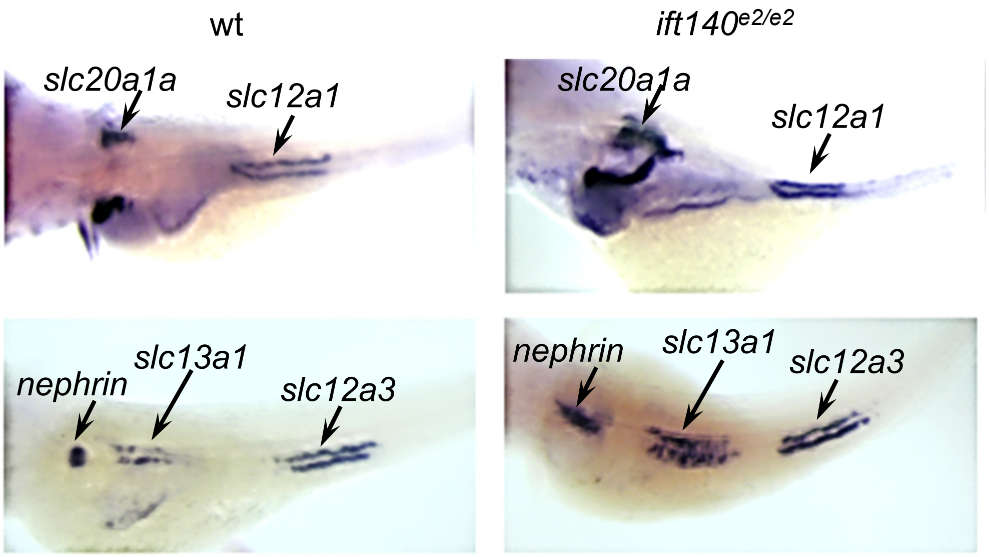


**Supplemental Figure 3. Inactivation of *ift140* does not affect kidney development**
Glomerular and tubular segmentation patterns are illustrated by *in situ* hybridization at 4 dpf. Markers include *nephrin* (podocyte marker), *slc20a1a* (proximal convoluted tubular marker), *slc13a1* (proximal straight tubular marker), *slc12a1* (distal early tubular marker), and *slc12a3* (distal late tubular marker). Extended *nephrin* expression and broader *slc13a1* expression indicate dilations in the glomerular-neck region and proximal tubular segment.

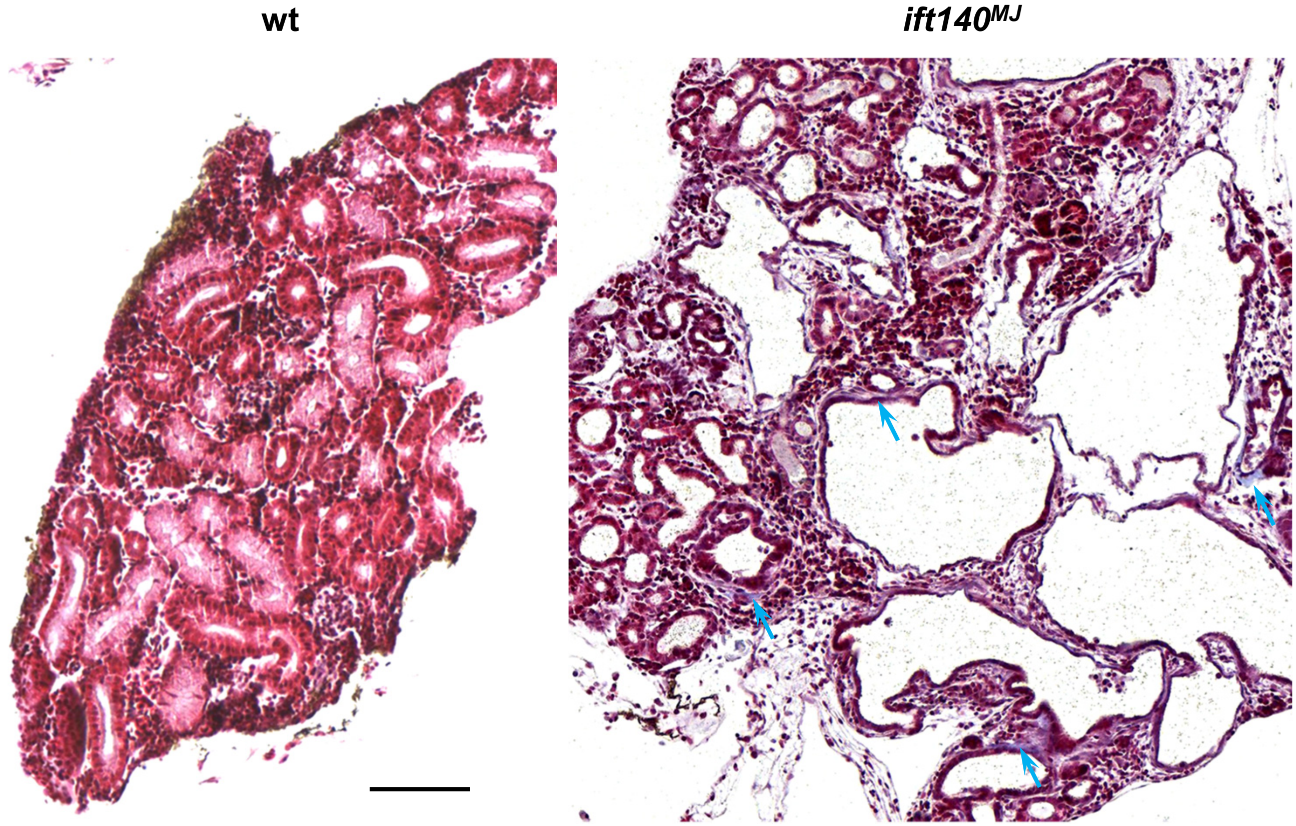


**Supplemental Figure 4. Fibrosis in the kidneys of adult *ift140$^{MJ}$* fish**

Masson's trichrome staining was performed on kidney tissues from adult zebrafish. Arrowheads indicate mild tubulointerstitial fibrosis observed in *ift140$^{MJ}$* fish. Scale bar: 100 µm.

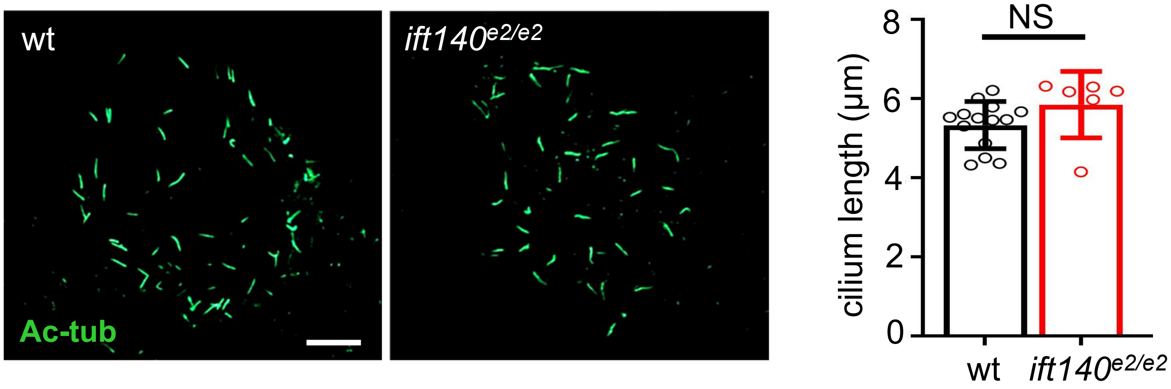


**Supplemental Figure 5. *ift140*$^{e2/e2}$ embryos display normal cilia in the Kupffer's vesicle**

Anti-acetylated tubulin antibody staining (Ac-tub) was performed on 10-somite stage embryos. Shown are cilia in the Kupffer's vesicle region and the quantification of cilium length. NS: not statistically significant (*P*>0.05). Scale bar: 20 μm.

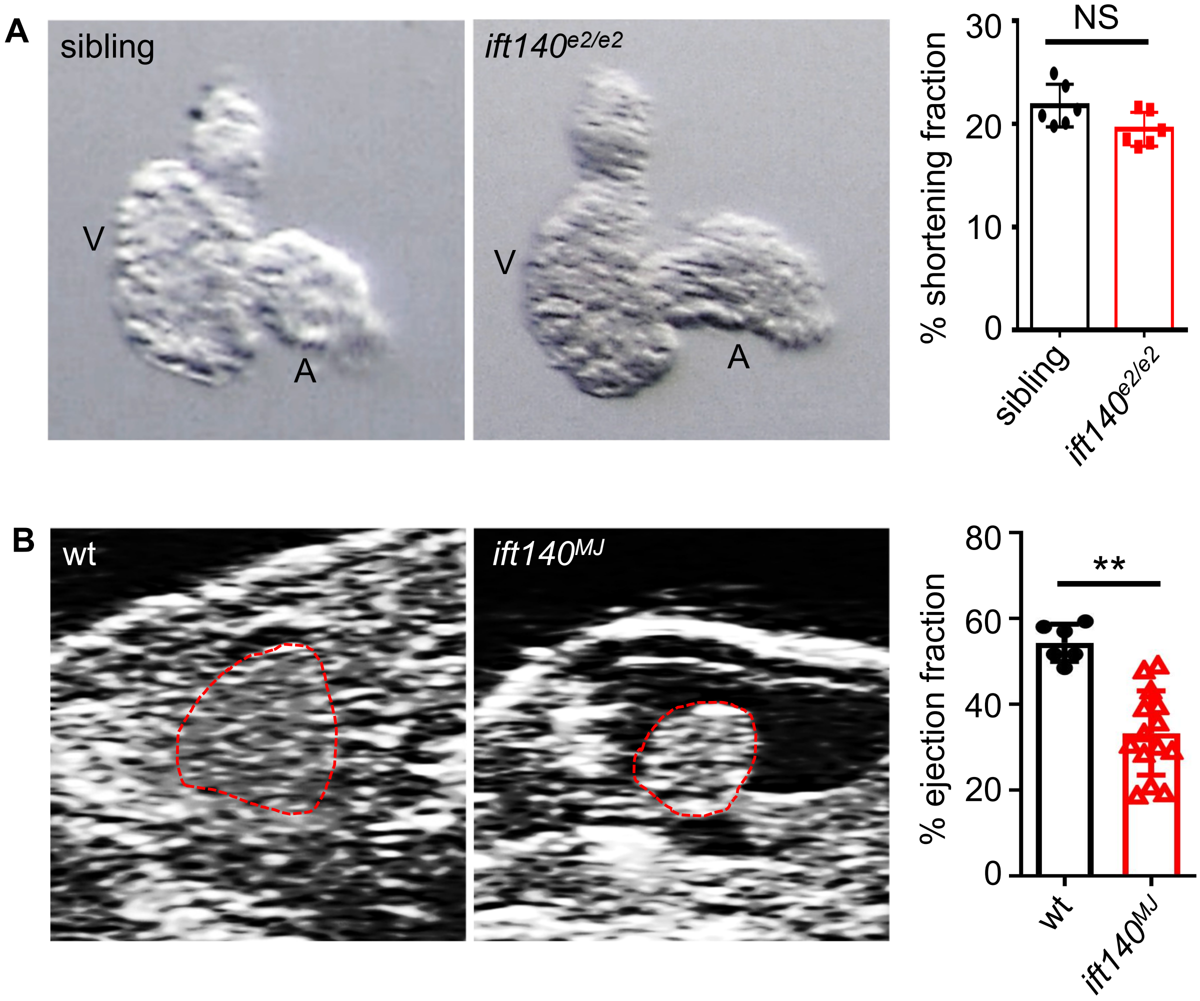


**Supplemental Figure 6. Cardiac phenotypes in *ift140*$^{e2/e2}$ embryos and adult *ift140*$^{MJ}$ fish**

(**A**) At 4 pdf, *ift140*$^{e2/e2}$ embryos exhibit generally normal cardiac morphology and function. Representative images of dissected hearts are shown, alongside quantitative analysis of cardiac function measured by the shortening fraction. (**B**) Adult *ift140*$^{MJ}$ fish at 3 months of age display defects in cardiac morphology and function. Representative echocardiography images extracted from movies of beating hearts are shown, accompanied by quantification of cardiac function through measurements of ejection fraction. **: $P$<0.01. NS: not statistically significant ($P$>0.05).

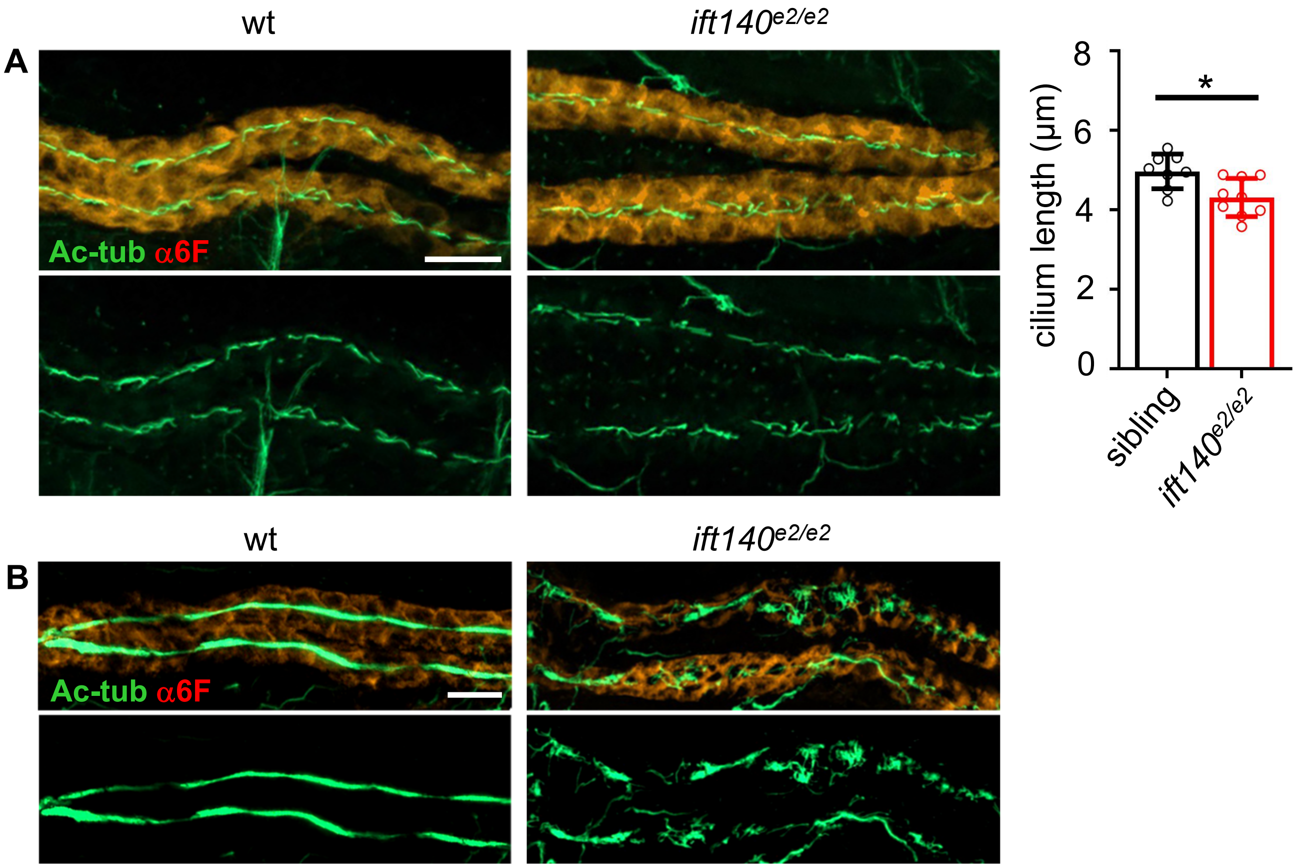


**Supplemental Figure 7. The effect of *ift140* inactivation on the length of distal single cilia and the orientation of multi-cilia bundles in the 2 dpf pronephros**
Whole-mount immunostaining was conducted on 2 dpf embryos. Cilia were visualized using an antibody against acetylated α-tubulin (Ac-tub), while the pronephros was labeled with an antibody against $Na^+/K^+$ ATPase (α6F). (**A**) Representative images of distal single cilia, along with quantification of cilium length. (**B**) Representative images of multi-cilia bundles. *: $P<0.05$. Scale bar: 20 μm.

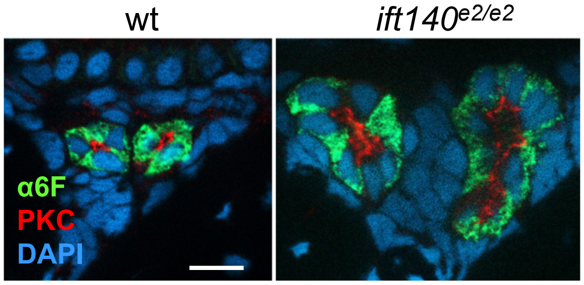


**Supplemental Figure 8. *ift140$^{e2/e2}$* embryos exhibit normal apico-basolateral polarity in kidney tubular epithelial cells**

Frozen sections of 5 dpf embryos were immunostained with antibodies against atypical PKC (red), marking the apical surface of kidney tubular epithelial cells, and $Na^{+}/K^{+}$ ATPase (α6F, green), marking the basolateral surface of kidney tubular epithelial cells. Scale bar: 10 μm.

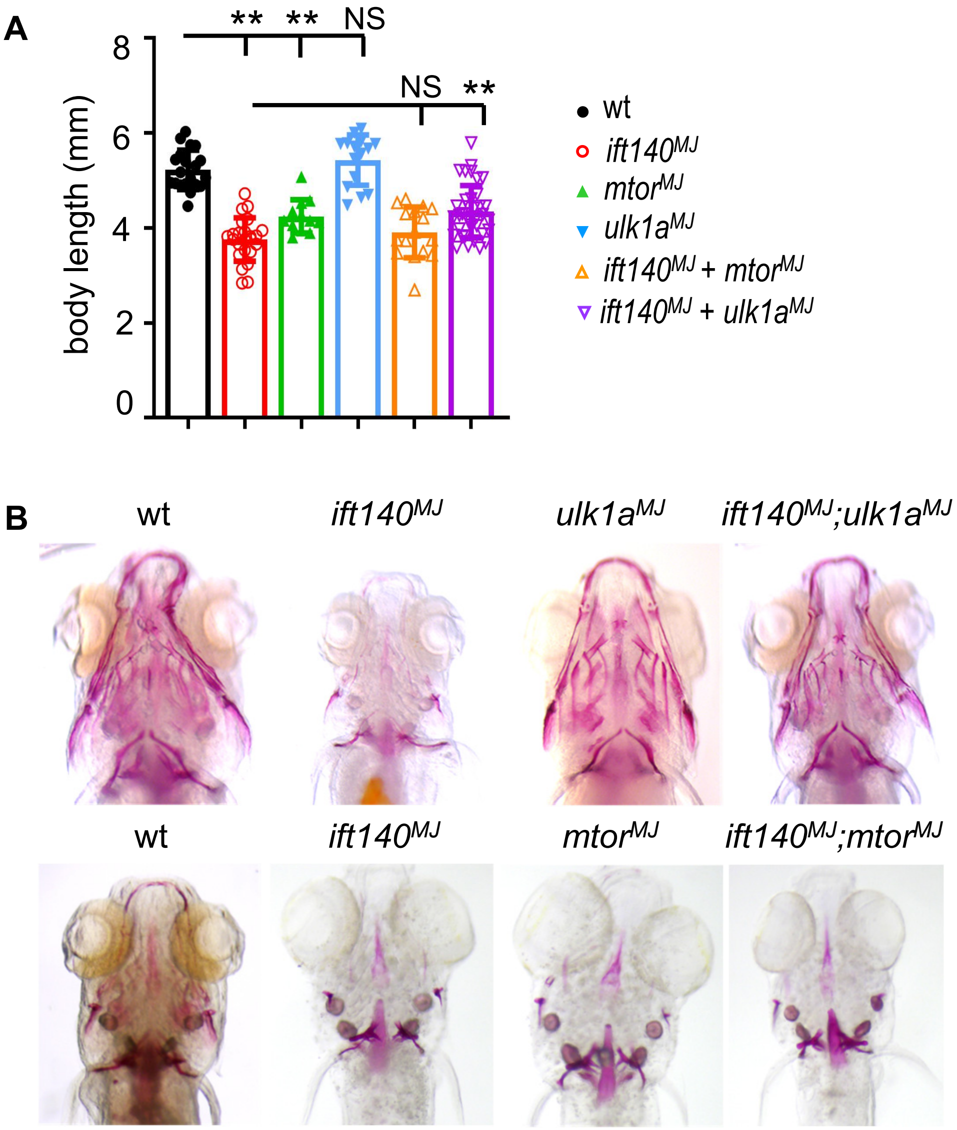


**Supplemental Figure 9. Effects of *ulk1a* and *mtor* inhibition on body size and calcified bone structures in *ift140$^{MJ}$* larvae**

(**A**) Knockdown of *ulk1a*, but not *mtor*, restored body length in *ift140$^{MJ}$* larvae. Embryos were injected with sgRNAs targeting *ift140, mtor*, or *ulk1a,* either individually or in combination. Body length were measured at 16 dpf. (**B**) Knockdown of *ulk1a*, but not *mtor*, promoted the development of calcified bone structures in *ift140$^{MJ}$* larvae. Alizarin Red staining was performed on sgRNA-injected larvae at 26 dpf (upper panel) and 16 dpf (lower panel). Ventral views of craniofacial bones are shown. **: *P*<0.01. NS: not statistically significant (*P*>0.05).

### Supplemental Table 1: List of the sequences for F0-based MMEJ screen and primers for quantifying knockout score

| Target Genes | sgRNA Sequence | Frame Shift | Strand | Deletion Sequence | PCR Sequence |
|---|---|---|---|---|---|
| *camk2a* | GCTGTGAAGTTGGCAGATTTTGG | Yes | forward | AGATTTTGGC | F: GCCAATGTCTGGTGGTACAG |
| | | | | | R: AAAATCTGAACACTGAAGGAGC |
| *camk2b1* | CAAGAACGCTGCTGTGAAGCTGG | Yes | forward | TGAAGCTG | F: AAATCATGTAGGAGCATCGC |
| | | | | | R: TGTGTTTAAGGCAGGAGAAATG |
| *nfatc1* | ATTCTCCTTGTGGCGGTAAACGG | Yes | forward | TAAACGG | F: TGCTTCTCCACGCCATTCTC |
| | | | | | R: AACCAACTGTCCTCCGTCAC |
| *mapk3* | GGAGGATTTTGATCTCTCTCAGG | Yes | complement | AGAG | F: CGCACTGTGAGTGTTTAAGG |
| | | | | | R: AACTATTTCTTTCCCTGCTGC |
| *map2k1* | CCCTACATCGTGGGCTTCTATGG | Yes | forward | CTATGGGGCTT | F: TTATGCAGTTGATCCACCTTG |
| | | | | | R: AGCCCATTTAACAGTAGTGTG |
| *mtor* | GAAAGAGAAAGGGATGAATAAGG | Yes | forward | ATAAGGATGA | F: AGGTGCAGCCATTCTTTGAT |
| | | | | | R: TCATACCTCTCCCTCCATAC |
| *pik3r1* | CAGACTGTCAAGTGTGGATGGGG | Yes | forward | ATGG | F: CACAGATGGGAAGAGTCTGTATG |
| | | | | | R: CATGTAGCCAAGTGCAAGTAATG |
| *ulk1a* | ATCCTTCTCTCATACAGCACAGG | Yes | forward | CACAG | F: CTGATCCATAACCAATCTGCG |
| | | | | | R: GTCTGAGTGAGGACACCATC |
| *tfeb* | TGGCTGGCGTGGGCTCAGGTGGG | Yes | complement | GCCCAC | F: AGTCAAACCCAGATACACAGGC |
| | | | | | R: TCATGCGGGACCAAATGCAG |
| *ulk2* | ATGATGCCAAAGCTGACCTGTGG | Yes | forward | ACCTG | F: CTGAAGGCTGTCTCAGATTG |
| | | | | | R: ACTTTAGGCATTCAACCGTAG |
| *atg5* | GTGCTTCGAGATGTTTGGTTTGG | Yes | forward | GTTTG | F: GAACAGATTGGTAGCTGAAGAC |
| | | | | | R: CTTAACTGTTTAACTCATGGCG |
| *atg7* | CTGTGCCTCCAGCGGAACGACGG | Yes | forward | AACGACGG | F: GATTGCGTTTCATGTGTCG |
| | | | | | R: TCTGCCACAAATGTTACTGATG |
| *notch3* | ACGCAAGTACCATGGTGGCATGG | Yes | complement | CCATG | F: TCACACAACTATACTCAGCTCTTC |
| | | | | | R: CAACACATTTGCCACCATTCT |
| *tln1* | ATGATCCAAAGAAGGGCATTTGG | Yes | forward | ATTTGGC | F: CCAGAAGGTCATGGAAGAGATG |
| | | | | | R: AGCCAATCCTGATGAAACTAGAG |
| *rab11fip3* | GAAGATGTGGAGACAGACAGCGG | Yes | forward | ACAG | F: AGGCATCAGTGCAATCAGC |
| | | | | | R: GTCGTCTCCATCTGTGAGTG |
| *nek8* | GGCTGAGAAGCTTGAGGACTTGG | Yes | complement | CCTCAAGC | F: GTATTGTTGTGTGTCAGGATCG |
| | | | | | R: TTAGTGCCTTGTCCTCCAG |

MMEJ: microhomology-mediated end joining; PCR: polymerase chain reaction
